# Stimulatory and inhibitory G-protein signaling relays drive cAMP accumulation for timely metamorphosis in the chordate *Ciona*

**DOI:** 10.1101/2024.04.09.588679

**Authors:** Akiko Hozumi, Nozomu M. Totsuka, Arata Onodera, Yanbin Wang, Mayuko Hamada, Akira Shiraishi, Honoo Satake, Takeo Horie, Kohji Hotta, Yasunori Sasakura

## Abstract

Larvae of the ascidian *Ciona* initiate metamorphosis tens of minutes after adhesion to a substratum via their adhesive organ. The gap between adhesion and metamorphosis initiation is suggested to ensure the rigidity of adhesion, allowing *Ciona* to maintain settlement after losing locomotive activity through metamorphosis. The mechanism producing the gap is unknown. Here, by combining gene functional analyses, pharmacological analyses, and live imaging, we propose that the gap represents the time required for sufficient cAMP accumulation to trigger metamorphosis. Not only the Gs pathway but also the Gi and Gq pathways are involved in the initiation of metamorphosis in the downstream signaling cascade of the neurotransmitter GABA, the known initiator of *Ciona* metamorphosis. The mutual crosstalk of stimulatory and inhibitory G-proteins functions as the accelerator and brake for cAMP production, ensuring the faithful initiation of metamorphosis at an appropriate time and in the right situation.

## Introduction

Metamorphosis is a widespread feature of development that allows animals to have different functions between larval and adult stages^1^. As larvae become adults, the shape and various characteristics of their physiology, gene expression, behavior, and lifestyle change. With these changes, animals can focus on a limited number of biological activities at each stage to increase feeding, growth, dispersal, and reproduction efficiencies. This biphasic mode of life is a widely conserved and ancient trait of animals, as evidenced by its presence in groups that maintain primitive states^2^, indicating that this long-conserved feature has contributed to animals’ flourishment. Therefore, to understand animal evolution, it is essential to elucidate the mechanisms underlying metamorphosis.

A key characteristic of metamorphosis is that both internal and external conditions determine its initiation. Sufficiently matured (or metamorphically competent^3^) larvae start metamorphosis only when they meet an appropriate external condition to be adults. Various organic and inorganic external stimuli suited to the lifestyle of adulthood trigger metamorphosis. Many marine invertebrates exhibit a benthic lifestyle at the adult stage^4^. Their planktonic larvae have an adhesive organ that secretes adhesives and adheres to a substratum. The cues associated with the adhesion, such as the physical contact with the substratum and a chemical from organisms surrounding the adherence site, can trigger their metamorphosis. Upon starting metamorphosis, their larvae lose locomotive organs and transition into benthic adult forms. Determining the timing of metamorphosis is important for benthic animals, particularly sessile ones, to ensure their future survival and reproduction, as they become unable to relocate after metamorphosis. Therefore, sessile animals are suspected to have elaborate mechanisms to start metamorphosis only when a firm adhesion is achieved at an appropriate place. Although several reports describe this kind of phenomenon^5–7^, the mechanism by which the external condition is turned into an internal mechanism to trigger metamorphosis at the right timing remains elusive.

Ascidians are marine invertebrate chordates that are the closest living relatives to vertebrates^8–11^. The chordate features of ascidians are best represented by their tadpole larval shape. Like vertebrate tadpoles, ascidian larvae swim by beating their tail through neuromuscular activities^12,13^. However, ascidians lose the tadpole shape during metamorphosis, and they exhibit a sessile lifestyle at the adult stage^14,15^. Ascidian metamorphosis comprises several key events, such as tail regression, body axis rotation, and adult organ growth^16^ (Figure 1A).

**Figure 1.**
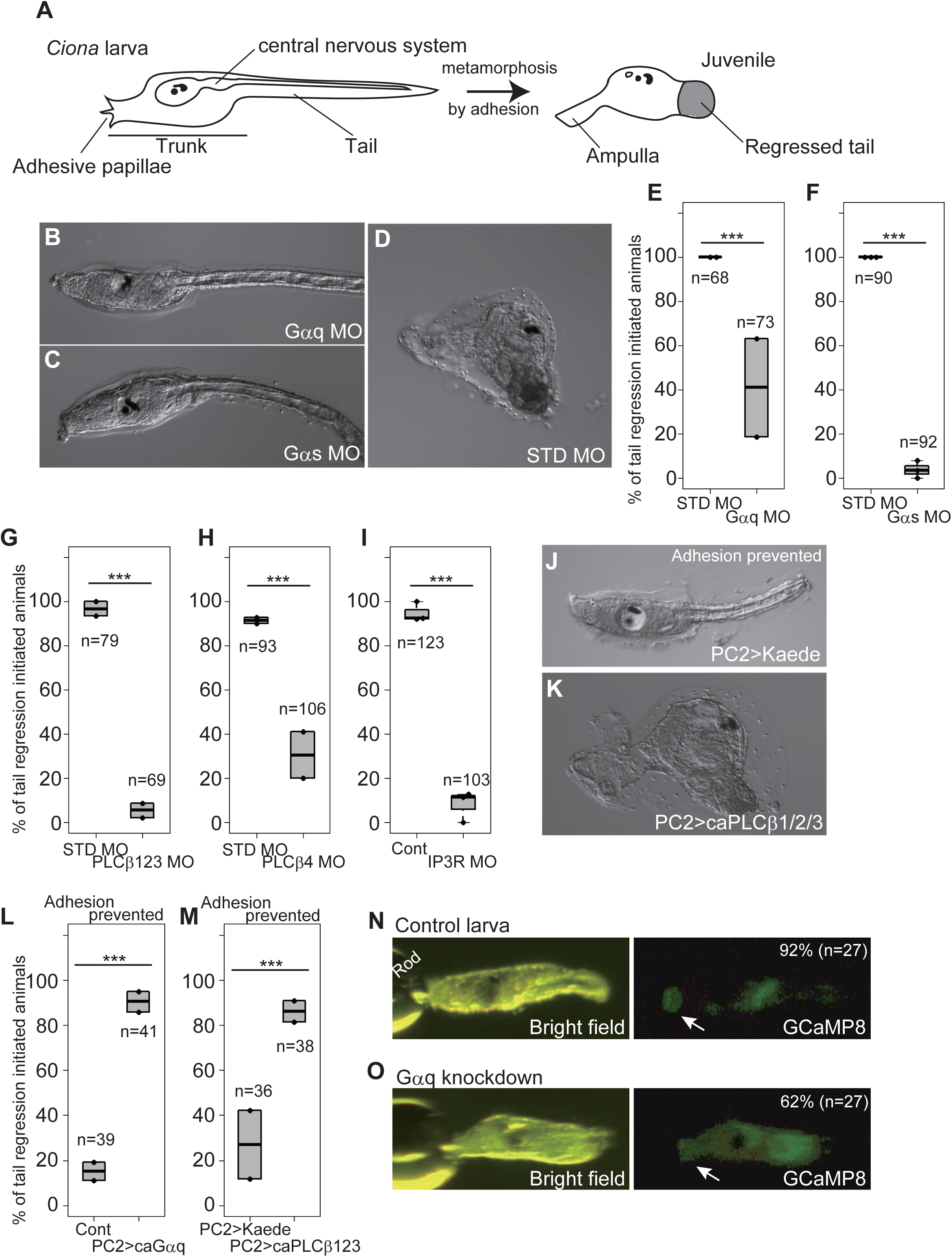
Gq and Gs pathways are required for *Ciona* metamorphosis. (A) Schematic illustration of *Ciona* metamorphosis. (B) A larva knocked down with *Gαq* using the antisense morpholino oligonucleotide (MO). Metamorphosis did not initiate at 2 days post-fertilization (2 dpf). (C) A *Gαs* knockdown larva. (D) A control animal injected with the standard (STD) MO. Metamorphosis initiated, as indicated by the completion of tail regression. (E) Effect of *Gαq* knockdown on the percentage of metamorphosis initiation (indicated by the initiation of tail regression), shown as box-and-whisker plots. Dots indicate experiment replicates. ***, p<0.001 (Fisher’s exact test). n, number of examined larvae in total. (F) Effect of *Gαs* knockdown. (G-I) Effects of *PLCβ1/2/3*, *PLCβ4*, and *IP3R* knockdowns. (J-M) Results of the experiments where adhesion was prevented by tail amputation and laying larvae on agar-coated plates. (J) A control larva overexpressing the *Kaede* reporter in the entire nervous system using the *PC2 cis* element. (K) A larva overexpressing constitutive active (ca)*PLCβ1/2/3*. (L) Effect of ca*Gαq* overexpression. (M) Effect of ca*PLCβ1/2/3* overexpression. (N-O) Live imaging of Ca^2+^ transient, monitored by injecting *GCaMP8* mRNA. (N) Snapshot of the control larva injected with *GCaMP8* mRNA. An increase in the fluorescence intensity in the adhesive papillae (arrow) was observed. Rod, the position of the glass rod used to stimulate papillae. The percentage and number exhibit the rate of animals showing Ca^2+^ transient in the papillae. (O) A larva injected with *Gαq* MO plus *GCaMP8* mRNA. An increase in the fluorescence intensity in the papillae (arrow) was not observed even though the brightness of the whole body was raised.

Adhesion to a substratum triggers ascidian metamorphosis^16^. Ascidian larvae have adhesive organs called adhesive papillae, usually triangular protrusions at the anterior end. Adhesive papillae secrete a lectin-type mucus substance that is suspected to aid in binding to the substratum^17^. Moreover, adhesive papillae serve as mechanical sensors^18^. Although cells in the papillae are polymodal, sensing both mechanical and chemical stimuli^19^, the mechanical stimulus can trigger metamorphosis without such a chemical^18,20^ under laboratory conditions. The papilla is a neuronal organ that includes epidermal neurons positive for vesicular glutamate transporter (VGLUT) and GABA, which innervate axons toward the sensory vesicle (larval brain)^21–24^. The papilla neurons are responsible for initiating metamorphosis^18,25–27^. Moreover, our recent study suggests the involvement of a mechanoreceptor channel (the TRP channel) in the responsiveness to mechanical stimuli^26^. However, a simple physical adhesion is not sufficient to trigger metamorphosis. In our study using the model ascidian *Ciona intestinalis* Type A (it has been proposed that this species be renamed *C. robusta*^28–30^, and hereafter we call it *Ciona*), continuous adhesion for about 30 min is necessary before starting metamorphosis^31^. When larvae detach before reaching the critical period, they must adhere for 30 min again for metamorphosis. Therefore, it is thought that ascidian larvae somehow sense the duration of adhesion and initiate metamorphosis only when adhesion is firm enough to be maintained for the required period. *Ciona* larvae continue tail beating during adhesion to push their body to the substratum. We showed that the strength of force generated by this swimming activity influences the timing of metamorphosis initiation^26^. In this phenomenon, abolishment of swimming activity elongates the time from settlement until metamorphosis is initiated. The mechanisms for measuring the duration of adhesion and the strength of the force generated by adhesion are unknown.

To elucidate the mechanisms triggering ascidian metamorphosis, the signaling pathways must be characterized. Many studies have discovered signaling molecules as possible inducers of ascidian metamorphosis^19,32–38^. These molecules include neurotransmitters, suggesting that transmission of excitation in the nervous system, starting from the adhesive papillae, is crucial for metamorphosis. Recently, our group reported that the inhibitory neurotransmitter GABA plays a pivotal role in initiating *Ciona* metamorphosis^39^. Knockout and knockdown of the genes encoding the enzyme synthesizing GABA called glutamate decarboxylase (GAD), vesicular inhibitory amino acid transporter (VIAAT or VGAT), and metabotropic GABA receptor (GABABR), resulted in the perturbation of metamorphosis. Moreover, GABA administration induces metamorphosis without adhesion. Our previous studies demonstrated that GABA induces the secretion of gonadotropin-releasing hormone (GnRH)^39,40^; however, it remains unknown how the inhibitory neurotransmitter activates neuronal functions for initiating metamorphosis.

In this study, we addressed the characterization of the downstream cascade stimulated by GABA. Because GABABR is a G-protein-coupled receptor (GPCR)^41^, we searched for trimeric G-proteins necessary for metamorphosis. These G-proteins are heterotrimers of an α β and γ subunit^42^. Upon ligand binding, GPCR exchanges GDP of the α subunit to GTP, then the GTP-bound α subunit and the βγ complex are released from the GPCR. Although both the GTP-bound α subunit and the βγ complex have activities, the α subunit mainly determines the reaction specific to the G-protein type. Gαs and Gαi respectively activate and inhibit cyclic adenosine monophosphate (cAMP) synthesis. In contrast, Gαq increases intracellular Ca^2+^ ion concentration by promoting its secretion from the endoplasmic reticulum. We found that three G-proteins are activated in the downstream GABA cascade, resulting in cAMP elevation. Because the signaling includes stimulating and inhibiting cAMP synthesis, *Ciona* initiates metamorphosis only when a sufficient quantity of cAMP is accumulated due to sustained adhesion. Our results revealed the ingenious mechanism that permits *Ciona* to start metamorphosis only when achieving an appropriate adhesion firm enough to relinquish swimming ability and commence sessile adult life.

## Results

### Characterization of G-proteins necessary for metamorphosis initiation

A previous study^43^ identified ten genes encoding Gα proteins from the *Ciona* genome, and we found nine gene models corresponding to these proteins in the latest genome assembly^44^ (Table S1). Gα proteins are classified into four major groups (Gαs, Gαi, Gαq, and Gα12/13), which have distinct functional properties^45^. *Ciona* has one gene encoding an unambiguous ortholog in each of the four groups (Figure S1A). Moreover, a previous study^43^ suggested the presence of another Gαq protein in the *Ciona* genome. Our phylogenetic tree suggested that Gαq (Gαq_Chr11) is orthologous to human GNA15 (Figure S1A).

The remaining four *Ciona* genes (their gene models are KY21.Chr2.875, KY21.Chr4.943, KY21.Chr9.455, and KY21.Chr8.580) encode divergent Gα proteins. The Gα protein encoded by KY21.Chr8.580 has truncation at its N-terminal part, and we omitted this protein from the phylogenetic analysis. Our phylogenetic analyses showed that the remaining divergent Gα proteins have a strong affinity to the Gαi/o family proteins (Figure S1A). Moreover, the amino acid residues at the C-terminal end of these proteins are more similar to human Gαi than the other Gα protein families (Figure S1B). The C-terminal residues, particularly glycine at position 3, are essential for determining the Gi partner of GPCR^46^. We tentatively name these proteins dvGαi_Chr2, dvGαi_Chr4, and dvGαi_Chr9. Gαi/o family proteins have inhibitory activity that suppresses cAMP production^47^.

To verify the possible contribution of G proteins to the initiation of metamorphosis, we quantified the expression level of the Gα genes in the adhesive papillae using RNA-seq surrounding this region (Figure S1C and Table S1). Typical *Gαq* and three *Gαi* genes exhibited more abundant expression in the papillae, followed by *Gαs* and *Gα12/13*. Therefore, these Gα proteins were chosen as candidates for the protein(s) regulating metamorphosis initiation. Among them, we prioritized examining the Gα proteins having an excitatory function (Gαq and Gαs) rather than an inhibitory role, since previous studies suggested that excitatory events such as Ca^2+^ transient and neuropeptide secretion occur when *Ciona* metamorphose^18,39^. The primary function of Gα12/13 is the activation of Rho-mediated actin dynamics. Although this regulation is essential for tail regression^48^, this event occurs after metamorphosis initiation. Therefore, we omitted this gene from further analysis in this study.

We knocked down the *Gαq* and *Gαs* genes using antisense morpholino oligonucleotides (MOs). We found that both these genes are necessary for metamorphosis. The morphants of these genes showed a reduced rate of metamorphosis initiation even though they were allowed to adhere to the base of culture dishes, which was sufficient for control larvae to initiate metamorphosis (Figure 1A-F). *Gαs* MO exhibited almost complete perturbation of metamorphosis, while the effect of *Gαq* MO was milder (Figure 1E and F).

Gαq activates phospholipase C beta (PLCβ) to produce inositol triphosphate (IP3)^49^. IP3 is received by its receptor on the endoplasmic reticulum (ER) that releases calcium ion (Ca^2+^). The *Ciona* genome encodes two PLCβ (PLCβ1/2/3, PLCβ4) and one IP3 receptor (IP3R) (Figure S2 and Table S1). We knocked down the *PLCβ1/2/3*, *PLCβ4*, and *IP3R* genes. The knockdown larvae of these three genes failed to start metamorphosis (Figure 1G-I). The effect of *PLCβ4* MO was weaker than those of the other MOs, suggesting that this PLC plays an auxiliary role. These results suggest that Gαq initiates metamorphosis by the conventional Ca^2+^ pathway mediated by PLCβ and IP3/IP3R. To confirm this, we overexpressed constitutively active forms of Gαq (caGαq)^50^ and of caPLCβ1/2/3^51^ throughout the entire nervous system with the *cis* element of the gene encoding prohormone convertase 2 (PC2)^52–54^. When the microinjected animals reached the larval stage, they were cultured on agar-coated dishes after tail amputation to prevent adhesion. In this condition, the control larvae rarely started metamorphosis because of the absence of adhesion (Figure 1J). In contrast, ca*Gαq*- or ca*PLCβ1/2/3*-overexpressed larvae initiated metamorphosis without adhesion (Figure 1K-M).

Adhesive papillae exhibit a Ca^2+^ increase soon after sensing an adhesive stimulus^18,26^. We examined whether this Ca^2+^ transient is dependent on the Gq pathway. Compared to controls, significantly fewer larvae injected with *Gαq* MO plus *GCaMP8* mRNA exhibited GCaMP8 fluorescence elevation in the adhesive papilla upon stimulation (Figure 1N, O). This result suggests that the Gq pathway is activated upon adhesion to cause a Ca^2+^ transient in the adhesive papilla.

### The Gs pathway initiates metamorphosis by activating cAMP synthesis

The involvement of Gs in metamorphosis was confirmed by the overexpression of a constitutively active form of Gαs (caGαs)^55^ in the nervous system. This overexpression resulted in the enhanced initiation of metamorphosis without adhesion (Figure 2A-C). The Gs pathway activates adenylate cyclase (AC) to produce cAMP^56^. We previously reported that cAMP can induce metamorphosis^40^. Because externally added cAMP is not a strong inducer of metamorphosis, we attempted to confirm this hypothesis through another experiment. Theophylline increases cAMP by inhibiting the cAMP-degrading enzyme phosphodiesterase (PDE)^57^. We treated wild-type larvae with theophylline after tail amputation, and we observed that most theophylline-treated larvae completed tail regression without adhesion (Figure 2D-F). Theophylline has several target proteins in addition to PDE^58^. To further confirm that cAMP is responsible for the initiation of metamorphosis, we overexpressed photo-activating AC (bPAC)^59^ in the nervous system.

**Figure 2.**
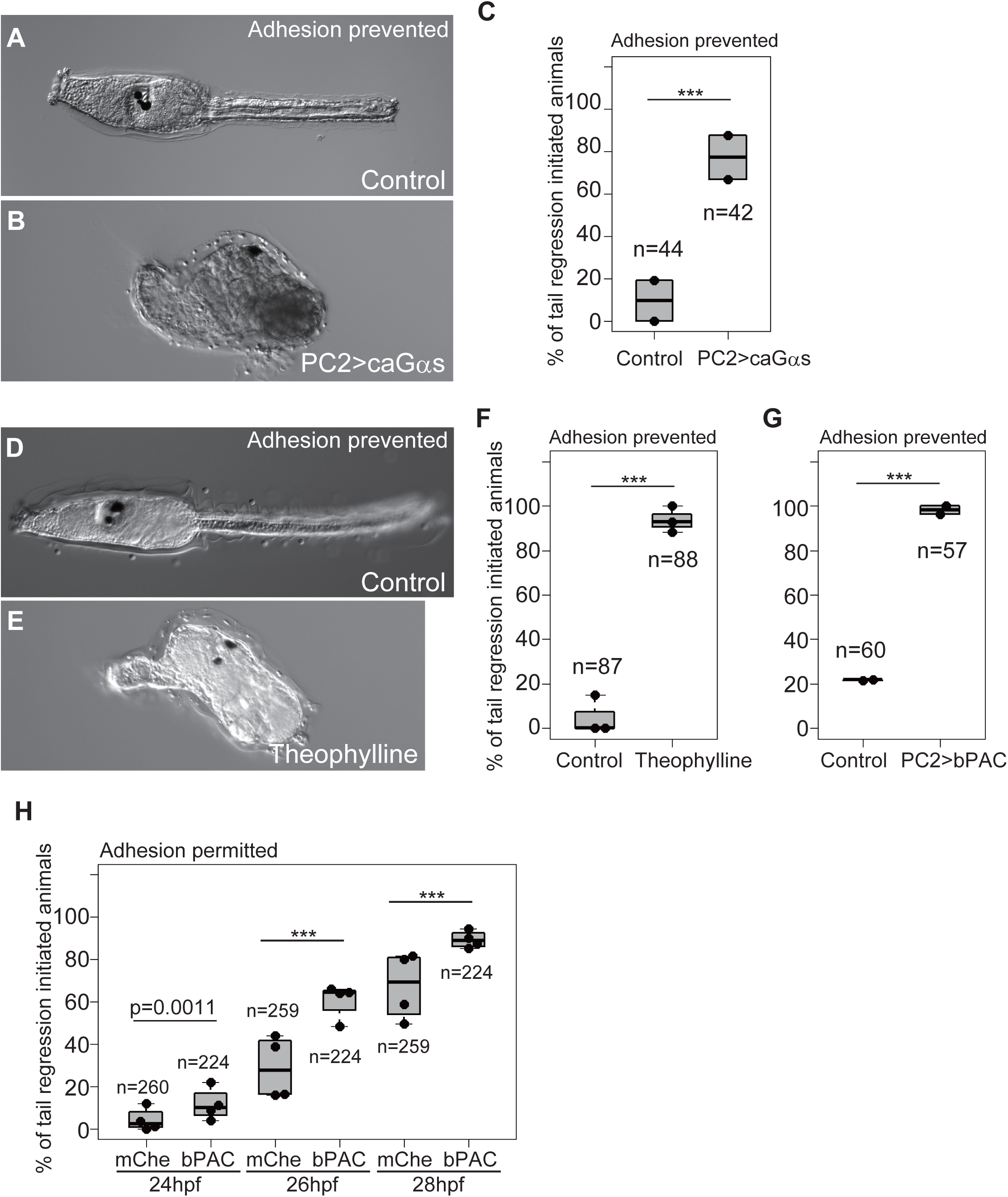
cAMP initiates *Ciona* metamorphosis. (A) An adhesion-prevented control (uninjected) larva at 2 dpf. Metamorphosis did not initiate. (B) An animal with adhesion prevention and overexpression of ca*Gαs* in the entire nervous system. Metamorphosis initiated. (C) Effect of ca*Gαs* overexpression. (D) A control (uninjected) larva. (E) A theophylline-treated animal. Metamorphosis initiated. (F) Effect of theophylline treatment. (G) Effect of *bPAC* overexpression in the nervous system. (H) Effect of *bPAC* overexpression over time. In this experiment, adhesion was permitted. Control groups (overexpressed with mCherry) exhibited a slower initiation of metamorphosis than bPAC-overexpressed groups.

The *bPAC*-overexpressed larvae regressed their tails without adhesion, suggesting that a cAMP increase triggers metamorphosis (Figure 2G).

Using bPAC, we addressed whether enhanced cAMP production facilitates the initiation of metamorphosis. At 24 hours post-fertilization (hpf), only a few animals in both the *bPAC*-overexpressed and control groups initiated metamorphosis because 24 hpf is somewhat too early for *Ciona* larvae to be metamorphically competent^31^ (Figure 2H). Even in this condition, the *bPAC*-overexpressed group exhibited a statistically higher rate of metamorphosis-initiated larvae. The proportion of animals initiating metamorphosis increased over successive time points, with the *bPAC*-overexpressed group consistently showing a statistically higher rate of metamorphosis compared to controls. These results support the idea that cAMP plays a role as a timer for *Ciona* metamorphosis; accumulation of this molecule to exceed a threshold could trigger metamorphosis. Because many of the *bPAC*-overexpressed larvae did not initiate metamorphosis at 24 hpf, similar to control larvae, cAMP accumulation does not alter the timing of acquiring metamorphic competence.

### Gq-Gs pathways work in the adhesive papillae for metamorphosis

The above results showed that activation of the Gq and Gs pathways are the key events in initiating metamorphosis. Gαq is necessary to induce Ca^2+^ transients in the adhesive papillae, suggesting that the Gq pathway functions in this region. If both Gq and Gs function in the papillae, they should be expressed in the adhesive papilla. The transcriptome analysis of the larval papilla region showed that the genes encoding Gq and Gs pathway proteins are expressed in this region (Figure S1C, Figure S3 and Table S1). Therefore, Gq and Gs could function in the papilla to initiate metamorphosis.

As shown above, theophylline induced metamorphosis without settlement. However, when the papillae were removed from larvae, the average rate of metamorphosis induction by theophylline was reduced from 100% to 30% (Figure 3A-C). This suggests that cAMP elevation in the adhesive papillae is essential for starting metamorphosis. The overexpression of ca*Gαq*, ca*PLCβ1/2/3*, ca*Gαs*, and *bPAC* by the *PC2 cis* element resulted in the initiation of metamorphosis without adhesion. Like theophylline, amputation of the papillae reduced their effects in starting metamorphosis (Figure 3D), confirming that activation of the Gq and Gs pathways in the adhesive papillae triggers metamorphosis. Among these experiments, ca*PLCβ1/2/3* overexpression was the most sensitive to papilla amputation, suggesting that PLCβ1/2/3 acts specifically in the papillae during metamorphosis.

**Figure 3.**
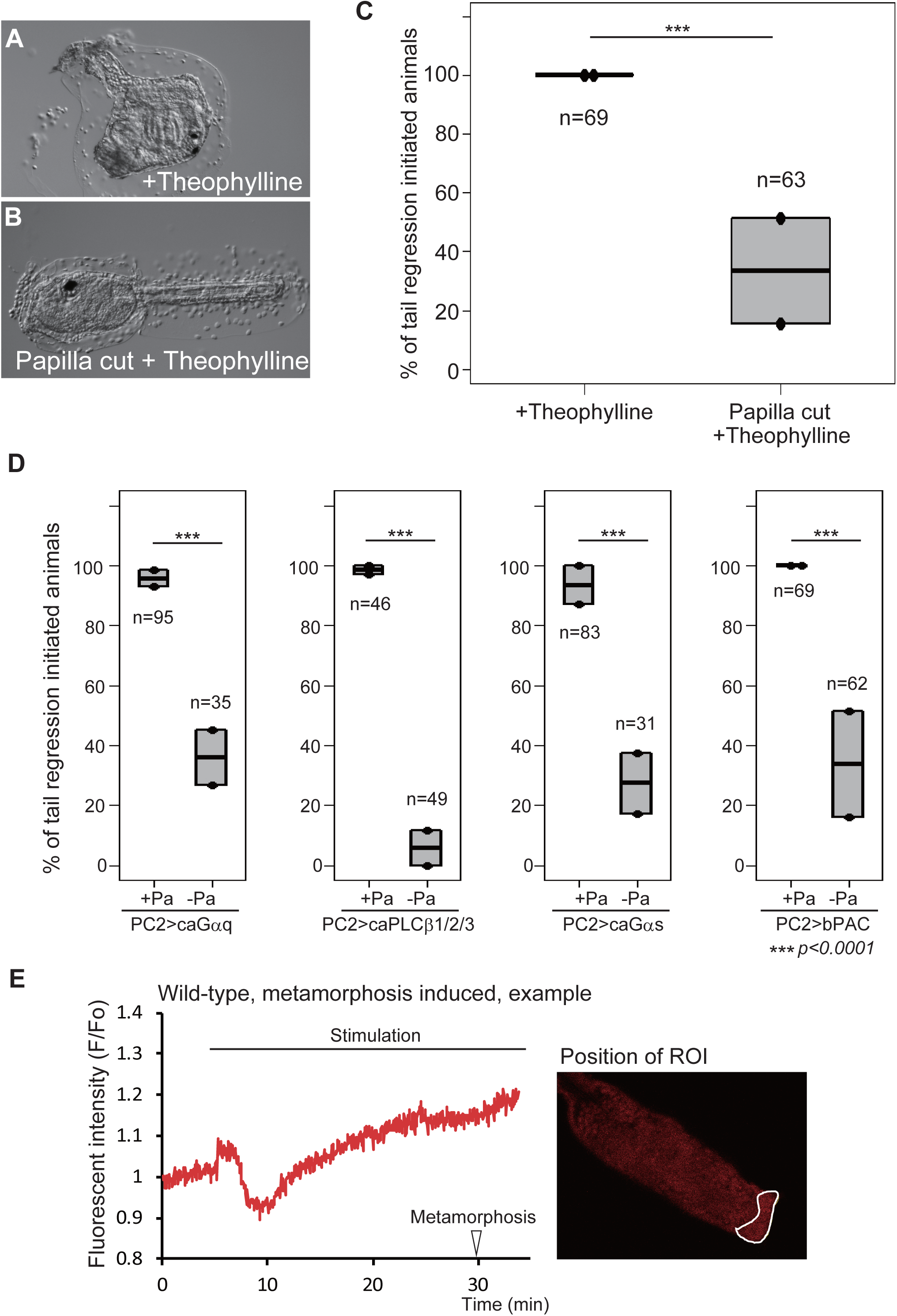
Gq and Gs pathways are activated in the adhesive papillae to initiate metamorphosis. (A) A theophylline-treated animal. Metamorphosis initiated. (B) A papilla-amputated larva treated with theophylline. Metamorphosis did not initiate. (C) Effect of papilla amputation on theophylline treatment. (D) Effects of papilla amputation on ca*Gαq*, ca*PLCβ1/2/3*, ca*Gαs*, and bPAC overexpression. (E) Measurement of cAMP concentration, as indicated by the fluorescence of Pink Flamindo. Left, the quantification of Pink Flamindo fluorescent intensity is shown for a larva that initiated metamorphosis at the time indicated by an arrowhead. The time of stimulation of the adhesive papillae with a glass rod is denoted by a bar. Right, the position of the region of interest (ROI). See also Figure S4.

If the Gs pathway is activated in the adhesive papillae, a cAMP increase should be observed in this region upon adhesion. We examined this possibility using a fluorescent cAMP indicator called Pink Flamindo (PF)^60^. After stimulation, the fluorescence of PF in the papillae temporarily decreased on average by 0.83-fold (n=5) from the initial intensity, followed by a gradual increase by 0.0102 per min, to reach 1.17-fold (Figure 3E and Figure S4A). The cAMP reduction and increase respectively started at 35 seconds and 4 min 40 seconds after stimulation on average. Such a reduction and increase in fluorescence intensity was not observed in the adhesive papillae of the larvae that had failed to initiate metamorphosis following stimulation (n=4; Figure S4B). Therefore, increased cAMP in the papillae serves to indicate that sufficient stimuli of adhesion have been received to induce metamorphosis. This strengthens our hypothesis that cAMP accumulation in the adhesive papillae determines the initiation of metamorphosis. Both the *Gαs* and *Gαq* knockdowns abolished the temporal cAMP decrease, and reduced the rate of subsequent accumulation (Figure S4C and D), indicating that these two events are dependent on the Gs and Gq pathways. However, neither knockdown abolished the increase in cAMP completely (as shown by the slope scores), suggesting the presence of a mechanism producing cAMP without their activations.

### GABA in the adhesive papillae is responsible for metamorphosis

Our previous study demonstrated that GABA is the chief neurotransmitter that induces metamorphosis^39^. To gain insight into the relationships between the GABA, Gq, and Gs pathways, we addressed whether GABA functions in the adhesive papillae for initiating metamorphosis, similar to Gq and Gs.

The previous studies and our transcriptome data suggest that *GAD* is expressed in the adhesive papillae^61^ (Table S1). Moreover, GABA-immunopositive signals have been detected in the papillae^23,61^. Therefore, papillae can provide GABA to stimulate themselves. Next, we examined whether the adhesive papillae can receive GABA to initiate metamorphosis. We showed that the metabotropic GABA receptor is responsible for the initiation of metamorphosis^39^. Using whole-mount *in situ* hybridization, the previous study did not detect the expression of two GABAB receptor (GABABR) genes in the papillae^61^; however, our transcriptome data detected the low-level expression of three genes encoding GABABR (Table S1) in the papillae, suggesting that adhesive papillae could receive GABA. GABA can induce metamorphosis without adhesion (Figure S5A)^39^. However, the amputation of adhesive papillae suppressed this activity (Figure S5B-C), suggesting that GABA reception by the papillae is responsible for starting metamorphosis.

The larval brain is the major domain expressing *GABABR*^61^. Therefore, it remains possible that GABA signaling in the brain stimulates the papillae in a retrograde manner to initiate metamorphosis. The connectome analyses of the larval nervous system did not suggest a nerve that inputs into the papillae from the brain^62,63^; however, retrograde stimulation would be possible through the volume transmission of a signaling molecule. To examine this possibility, we isolated larval fragments anterior to the brain (Figure S5D), followed by the administration of GABA and theophylline. Through the application of these chemicals, anterior fragments exhibited increased clarity, elongation, and retraction of papillae, which are the signatures of metamorphosis (Figure S5E-G). Because their responses to GABA were weaker than those observed in the theophylline treatment, we further examined whether the anterior fragments can respond to GABA by monitoring the expression of GABA-responsive genes. By comparing expression levels between control and GABA-administered larvae, we made a list of the genes upregulated by GABA in the papillae (Table S2). Among them, we compared the expression levels of two genes (KY21.Chr5.240 and KY21.Chr8.489) between GABA-administered and control anterior fragments. GABA-treated anterior fragments expressed these genes at levels at least three times higher than controls (Figure S5H). We concluded that GABA is secreted from and stimulates the papillae to start metamorphosis.

### Gs/cAMP is downstream of the Gq pathway

Our next question is: What are the upstream or downstream relationships between GABA, Gq/Ca2+, and Gs/cAMP in the adhesive papillae? We first examined the relationships between the Gq and Gs pathways. Theophylline ameliorated metamorphosis-failed phenotypes in *Gαq*, *PLCβ*, and *IP3R* knockdowns (Figure 4A-E). Moreover, ca*Gαs* overexpression in the nervous system significantly, although not completely, ameliorated the effect of *Gαq* MO (Figure 4F). These results suggest that the Gq pathway is upstream of the Gs/cAMP pathway. *Gαs* knockdown larvae exhibited Ca^2+^ transients in the adhesive papillae upon stimulation (Figure 4G). Because this Ca^2+^ transient is Gq-dependent (Figure 1O), this result confirms that the Gs pathway is downstream of the Gq-dependent Ca^2+^ increase.

**Figure 4.**
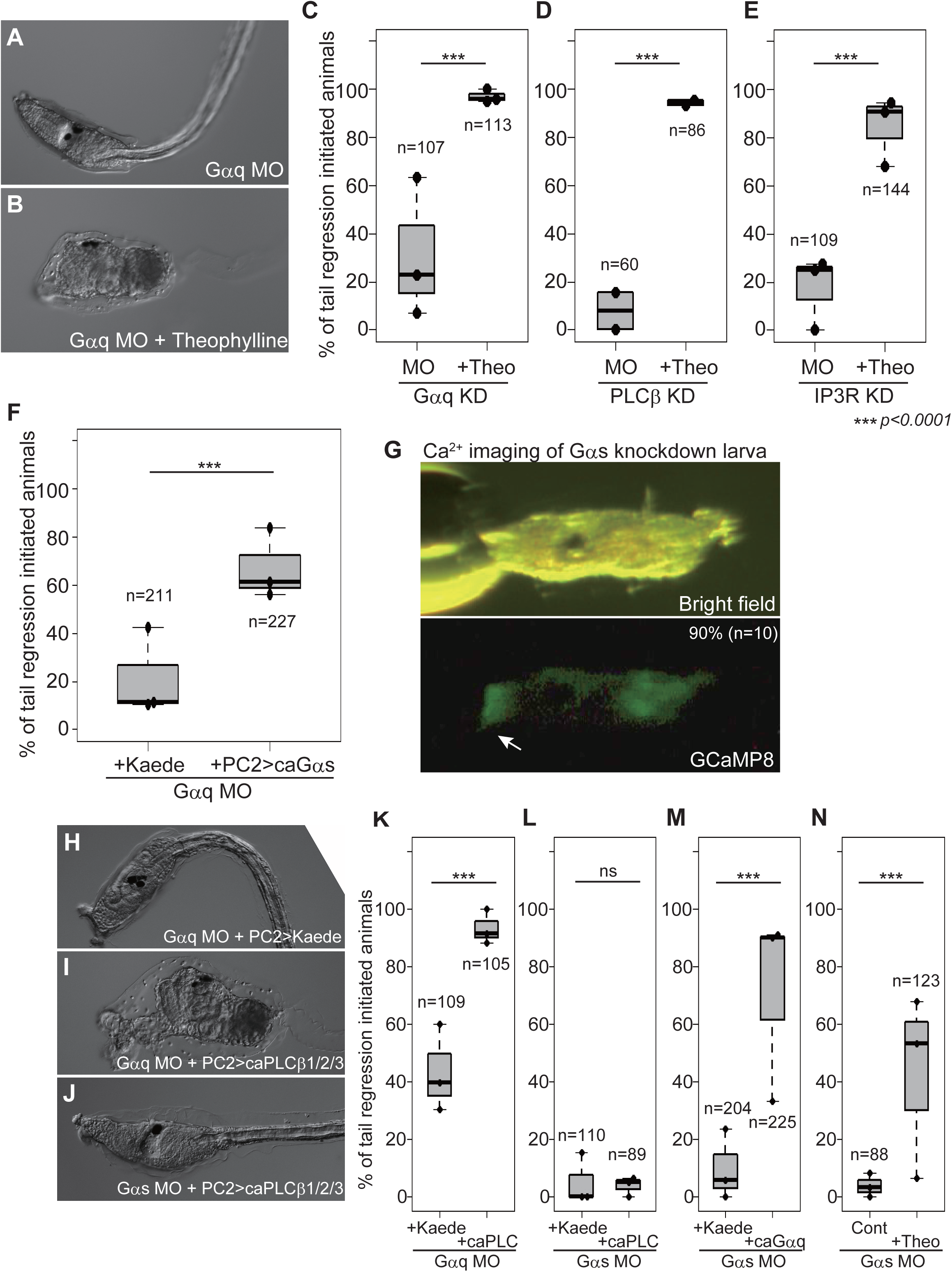
Relationships between Gq and Gs pathways in metamorphosis. (A) A *Gαq* knockdown larva. (B) Theophylline ameliorated the effect of *Gαq* knockdown. (C) Effect of theophylline on *Gαq* knockdown. (D) Effect of theophylline on *PLCβ* knockdown. In this experiment, *PLCβ1/2/3* and *PLCβ4* were knocked down simultaneously. (E) Effect of theophylline on *IP3R* knockdown. (F) Effect of ca*Gαs* overexpression on *Gαq* knockdown. (G) *Gαs* is not necessary for Ca^2+^ transient in the adhesive papillae (arrow) upon adhesion. The percentage and number represent the rate of animals showing Ca^2+^ transient in the papillae. (H) A *Gαq* knockdown and *Kaede*-overexpressed larva. (I) A *Gαq* knockdown and ca*PLCβ1/2/3*-overexpressed larva. (J) A *Gαs* knockdown and ca*PLCβ1/2/3*-overexpressed larva. (K) Effect of ca*PLCβ1/2/3* overexpression on *Gαq* knockdown. (L) Effect of ca*PLCβ1/2/3* overexpression on *Gαs* knockdown. (M) Effect of ca*Gαq* overexpression on *Gαs* knockdown. (N) Effect of theophylline treatment on *Gαs* knockdown.

To gain further insight into the epistatic order of the Gq and Gs pathways, we overexpressed constitutive active forms of Gq pathway proteins in *Gαs* morphants with the *PC2 cis* element. If Gq is upstream of the Gs pathway, forced activation of the Gq pathway by caGαq or caPLCβ1/2/3 would not ameliorate the *Gαs* MO effect. Indeed, ca*PLCβ1/2/3*, which rescued *Gαq* morphants well, failed to rescue *Gαs* morphants (Figure 4H-L). In contrast, ca*Gαq* significantly ameliorated the metamorphosis-failed phenocopies of *Gαs* morphants (Figure 4M), contradicting the hypothesis that Gq is upstream of the Gs pathway. One possibility is that the Gq pathway stimulates cAMP synthesis through a massive Ca^2+^ increase and protein kinase C (PKC) activation^64,65^. Indeed, *Ciona* larvae can synthesize cAMP without Gs, as suggested by the imaging of Pink Flamindo (Figure S4C), and by the partial rescue of *Gαs* morphants by theophylline treatment (Figure 4N). These data suggest that Gαs-independent AC could be a target of the Gq pathway. We concluded that the Gq pathway is upstream of the Gs pathway in the signaling cascade initiating *Ciona* metamorphosis.

### Interaction between GABA and Gq pathways

We next investigated the relationships between the GABA and Gq/Gs pathways. As our previous study showed, GABA pathway knockdown by *GAD* or *GABABR1* MO disrupted the initiation of metamorphosis (Figure 5A)^39^. Overexpression of ca*PLCβ1/2/3* and theophylline treatment ameliorated the metamorphosis-failed phenocopies of *GAD*/*GABABR1* morphants (Figure 5B-D). These results suggest that the GABA pathway is upstream of the Gq and Gs pathways. We observed Ca^2+^ transients in the adhesive papillae of *GAD* knockdown larvae. *GAD* morphants rarely exhibited a Ca^2+^ increase after stimulation (Figure 5E). Because the Ca^2+^ transient in the papillae is Gq-dependent, this result confirms that the GABA pathway is upstream of, or at least in parallel with, the Gq pathway.

**Figure 5.**
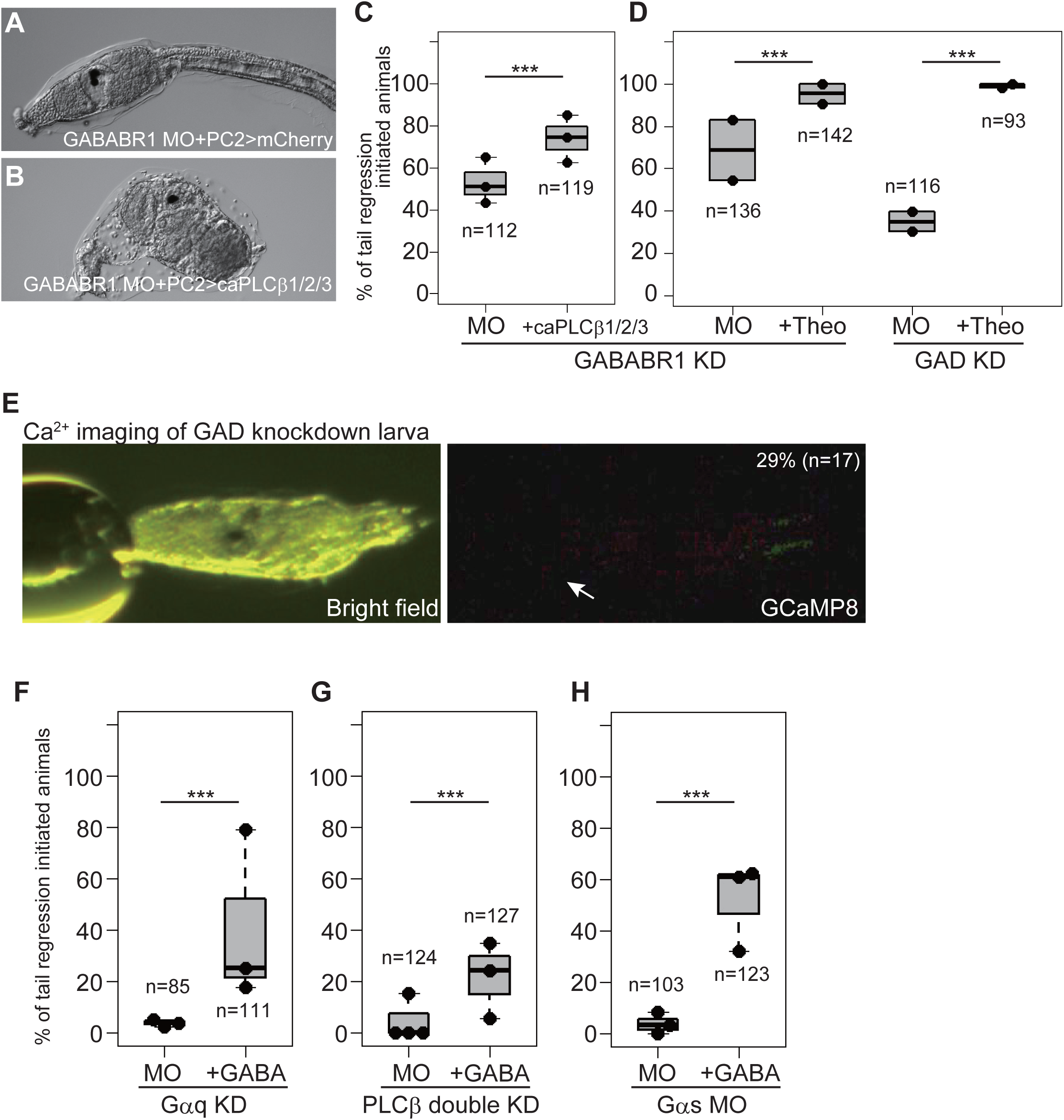
GABA functions in the adhesive papillae to induce metamorphosis. (A) A *GABABR1* knockdown and *mCherry*-overexpressed larva. (B) A *GABABR1* knockdown and ca*PLCβ1/2/3*-overexpressed larva. Metamorphosis initiated. (C) Effect of ca*PLCβ1/2/3* overexpression on *GABABR1* knockdown. (D) Effect of theophylline on *GABABR1* and *GAD* knockdowns. (E) GABA is necessary for Ca^2+^ transient in the adhesive papillae (arrow). The percentage and number represent the rate of animals showing Ca^2+^ transient in the papillae. (F) Effect of GABA on *Gαq* knockdown. (G) Effect of GABA on *PLCβ* knockdown. (H) Effect of GABA on *Gαs* knockdown. See also Figure S5.

However, puzzling results were obtained when we administered *Gαq*- and *Gαs*-knockdown larvae with GABA. If GABA is upstream of the Gq and Gs pathways, this chemical does not induce metamorphosis when the Gq or Gs pathway is disrupted. However, GABA partially but significantly ameliorated the metamorphosis-failed phenocopies of *Gαq*, *PLCβ*, and *Gαs* morphants (Figure 5F-H). Among the three morphants, GABA achieved more effective rescue in *Gas* knockdowns than *Gαq* or *PLCβ*. These results could be explained by assuming enhancement of the Gq pathway by GABA through PLCβ and another GABA-mediated metamorphic pathway bypassing Gq components. These possibilities are examined in the next section.

### Contribution of Gi to metamorphosis

The function of GABA as the potential upstream factor of Gq could be easily explained if GABABR is coupled with Gq. Although a few studies suggest this possibility^66^, Gi is regarded as the major G-protein coupled with GABABR^67^. A previous study demonstrated that the gene encoding typical *Ciona* Gαi is strongly expressed throughout the entire larval nervous system, including in adhesive papillae^68^, and our RNA-seq confirmed this (Figure S1C and Table S1). The knockdown of this gene (*Gαi*) exhibited a moderate (although statistically significant) reduction of metamorphosis rate (Figure S6A), suggesting the presence of another Gαi regulating metamorphosis.

Indeed, the *dvGαi_Chr2* knockdown larvae failed to initiate metamorphosis (Figure 6A-C), and this phenocopy was rescued by theophylline and GABA (Figure 6D), suggesting that *dvGαi_Chr2* is chiefly necessary for metamorphosis. The knockdown of *dvGαi_Chr4* resulted in a slight reduction of the metamorphosis rate (Figure S6B), suggesting the supportive role of this Gαi in the metamorphosis initiation. These data strengthen the hypothesis that GABABR regulates metamorphosis through Gi. The rescue of *dvGαi_Chr2* morphants by GABA might be attributable to a redundant function of the typical Gαi and dvGαi_Chr4 proteins in the papillae; however, we could not fully address this hypothesis due to a technical limitation. That is, the simultaneous knockdown of *dvGαi_Chr2* and *dvGαi_Chr4* did not suppress the amelioration of metamorphosis by GABA (Figure S6C). This might have been due to the presence of typical *Gαi*. The simultaneous knockdown of *dvGαi_Chr2* and typical *Gai* resulted in malformation of the body (Figure S6D), and we could not examine their responsiveness to GABA.

**Figure 6.**
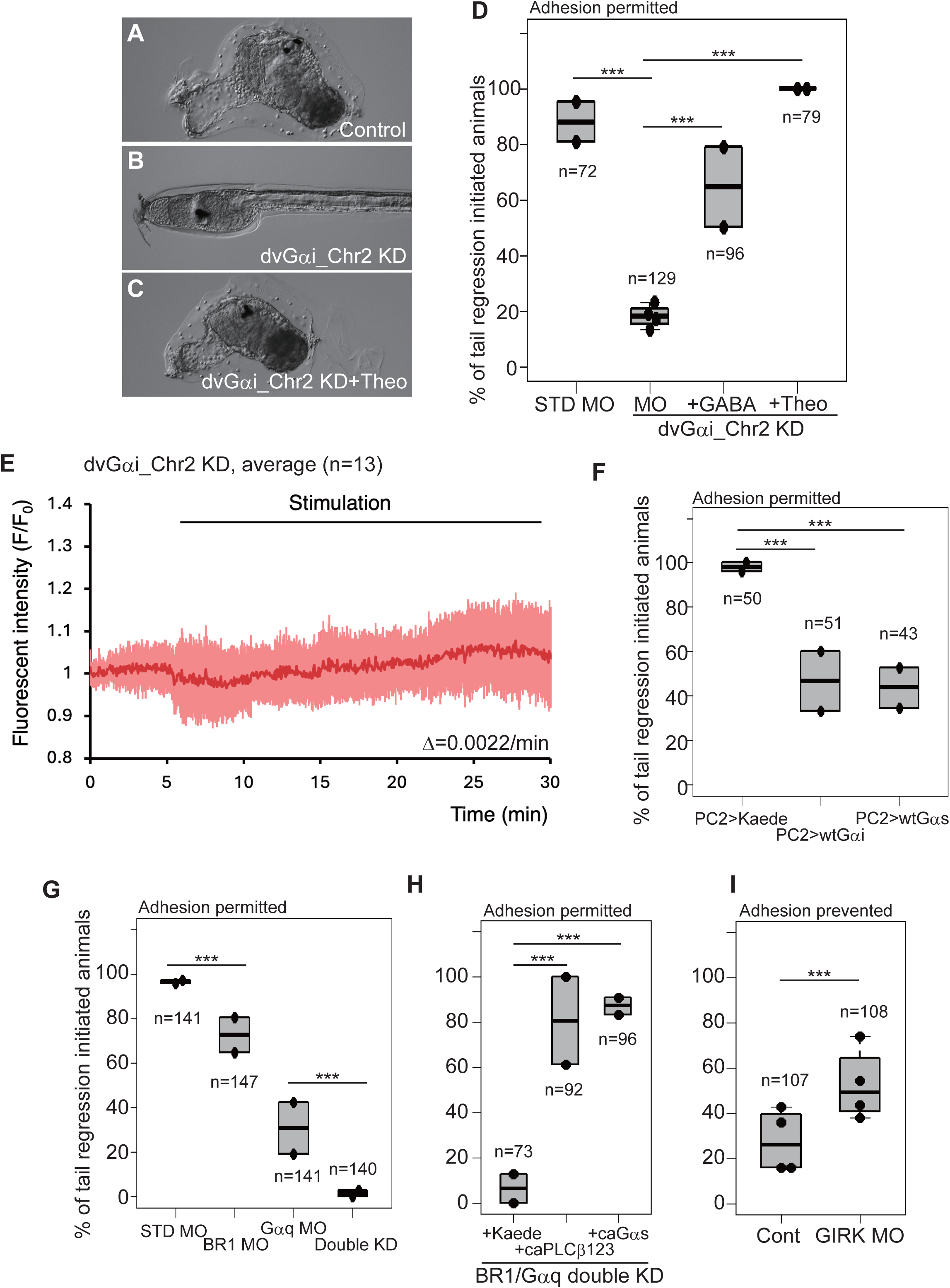
Gi is required for metamorphosis initiation. (A) A control animal injected with STD MO. (B) A *dvGαi_Chr2* knockdown larva. (C) A *dvGαi_Chr2* knockdown and theophylline-treated animal. (D) Effects of GABA and theophylline on *dvGαi_Chr2* knockdown. (E) Measurement of cAMP concentration in the *dvGαi_Chr2* knockdown larvae. The red and pink graphs represent the averaged Pink Flamindo fluorescent intensity and the standard deviations, respectively. The number in the parentheses indicates the number of examined larvae. The red graph is approximated to the linear graph to calculate its slope, which is shown as Δ. See also Figure S4. (F) Effects of wt*Gαi* and wt*Gαs* overexpression. (G) Effect of *GABABR1* and *Gαq* simultaneous knockdown. (H) Effects of ca*PLCβ1/2/3* and ca*Gαs* overexpressions on *GABABR1* and *Gαq* simultaneous knockdown. (I) *GIRK* knockdown promoted metamorphosis initiation. See also Figure S6.

We further explored the involvement of Gi upon metamorphosis as the upstream factor of the Gq-Gs pathways. Knockdown of *dvGαi_Chr2* resulted in the abolishment of temporal cAMP decrease and its subsequent accumulation upon adhesion (Figure 6E). Because these cAMP fluctuations are Gq- and Gs-dependent (Figure S4C-D), their dependence on dvGai_Chr2 supports that Gi is upstream of the Gq-Gs pathways. Gi is known to activate PLCβ through the Gβγi complex^69,70^.

Released βγ is inactivated by overexpressing normal (usually GDP-bound) Gα subunits because GDP-Gα quenches the βγ complex^71^. Overexpressing wild-type *Gαi* and *Gαs* in the nervous system significantly suppressed metamorphosis (Figure 6F), suggesting that the activated βγ complex is necessary for initiating metamorphosis. To confirm that the wild-type Gαi exerts its effect through sequestration of the βγ complex, we overexpressed a dominant-negative form of Gαi (dnGαi)^72,73^ which has reduced GDP/GTP affinity while maintaining βγ binding activity. dnGαi significantly reduced the occurrence of metamorphosis (Figure S6E), suggesting that the negative effect of Gαi on metamorphosis occurs through interaction with the βγ complex.

If Gβγ-mediated PLCβ activation is used in *Ciona* metamorphosis, PLCβ receives two independent inputs (Gαq and Gβγi) for its activation. These pathways could compensate for each other. We noticed that the MOs for GABA pathway genes and the *Gαq* MO did not disrupt metamorphosis as strongly as the *Gαs* MO (Figure 1E-F; 5C-D). The compensatory role could explain this phenomenon. Indeed, the simultaneous knockdown of *GABABR1* and *Gαq* resulted in a strong impairment of metamorphosis (Figure 6G), and these morphants initiated metamorphosis by overexpression of ca*PLCβ1/2/3* or ca*Gαs* (Figure 6H).

If the role of the GABA/Gi pathway relies specifically on PLCβ activation in the mechanism of metamorphosis, GABA administration would not rescue *PLCβ* morphants. However, GABA weakly but significantly induced metamorphosis in *PLCβ1/2/3* plus *PLCβ4* double-knockdown larvae (Figure 5G). Therefore, GABA/Gi is likely to activate another pathway that bypasses PLCβ. Gβγi is known to activate group III AC^64^. Its *Ciona* counterpart (AC5/6) is expressed in the adhesive papillae (Figure S3B), which could explain the results of the rescue experiments. In addition, there are two other major targets of Gβγi. One is the G-protein-activated inwardly rectifying potassium (GIRK) channel^74^. Our RNA-seq data on the papillae indicated the expression of two GIRK channel genes (Table S2). The GIRK channel negatively regulates the excitation of neurons through hyperpolarization. If the metamorphosis of *Ciona* is induced by the excitation of papilla neurons as suggested previously^18,26^, the GIRK channel is likely to regulate metamorphosis negatively. Indeed, the knockdown of one GIRK channel gene weakly enhanced the initiation of metamorphosis without settlement (Figure 6I). Although the negative role of the GIRK channel supports the involvement of Gβγi in the pathway of metamorphosis, this does not explain the PLCβ-independent activation of the metamorphic pathway by GABA/Gi.

The third function of Gβγi is to activate MAPK signaling through positive regulation of MEK^75^. PLCβ does not mediate this pathway. It has been suggested that MAPK signaling is required to induce metamorphosis^76,77^. To show that MEK activation is necessary for inducing metamorphosis mediated by GABA, we treated adhesion-prevented larvae with GABA plus U0126, a potent MEK1/2 inhibitor repeatedly used in ascidians^78,79^. U0126 significantly reduced the rate of metamorphosis induction by GABA (Figure S6F).

cAMP is also known to activate the pathway that involves MEK1/2^80^. We found that U0126 antagonized the effect of theophylline on metamorphosis (Figure S6G). Therefore, GABA and cAMP have the same target (MEK1/2) to initiate metamorphosis, suggesting their compensatory function and/or that cAMP synthesis is upregulated by GABA-Gβγi. These bypassing pathways could explain the amelioration of the phenocopies of *PLCβ* and *Gαs* morphants by GABA (Figure 5G and H).

### The constitutive function of wild-type Gαq

We found that *Ciona* wild-type Gαq (wtGαq) has constitutive activity. In contrast to wt*Gαs* and wt*Gαi*, overexpression of wt*Gαq* did not arrest metamorphosis (Figure S7A). Rather, its overexpression induced metamorphosis without settlement (Figure S7B). The constitutive wt*Gαq* activity was confirmed by the significant rescue of *Gαs* morphants (Figure S7C). As mentioned above, caGαq also has these activities (Figure 1L). ca*Gαq*-overexpressed larvae had a somewhat rounder trunk shape, abnormal tail bending, and frequent tail twitching, perhaps due to a constitutive increase in the cytosolic Ca^2+^ concentration (Figure S7D). Overexpression of wt*Gαq* did not show such abnormalities (Figure S7E), suggesting that wtGαq has milder activity than, or works through a different mechanism from, caGαq. To gain further insight into the constitutive effect of wtGαq, we constructed a dominant-negative form of Gαq^72^ that has reduced GDP/GTP binding activity while maintaining βγ binding activity. Overexpression of dn*Gαq* weakly inhibited metamorphosis (Figure S7F), suggesting that GDP/GTP exchange is important for the constitutive activity of wtGαq.

## Discussion

Among chordates, the tunicate ascidian is the only group that exhibits a sessile lifestyle at the adult stage. Understanding how ascidians acquired the mechanism to metamorphose into sessile adults is important for elucidating the evolution of chordates^81,82^. Through extensive molecular, physiological, and pharmacological analyses, we identified signaling molecules responsible for the initiation of metamorphosis of the ascidian *Ciona*. Combining the results with previous knowledge, we described a schematic that best represents the cascades running in the adhesive papillae upon adhesion to initiate metamorphosis (Figure 7). This working hypothesis will be the basis for future research on ascidian/tunicate metamorphosis. The hypothesis explains the characteristics of *Ciona* metamorphosis, the mechanisms of which have not yet been elucidated. The genes, proteins, and signaling pathways in this schematic are essential targets for elucidating how the ancestor of ascidians acquired the metamorphosis system peculiar to this group during evolution.

**Figure 7.**
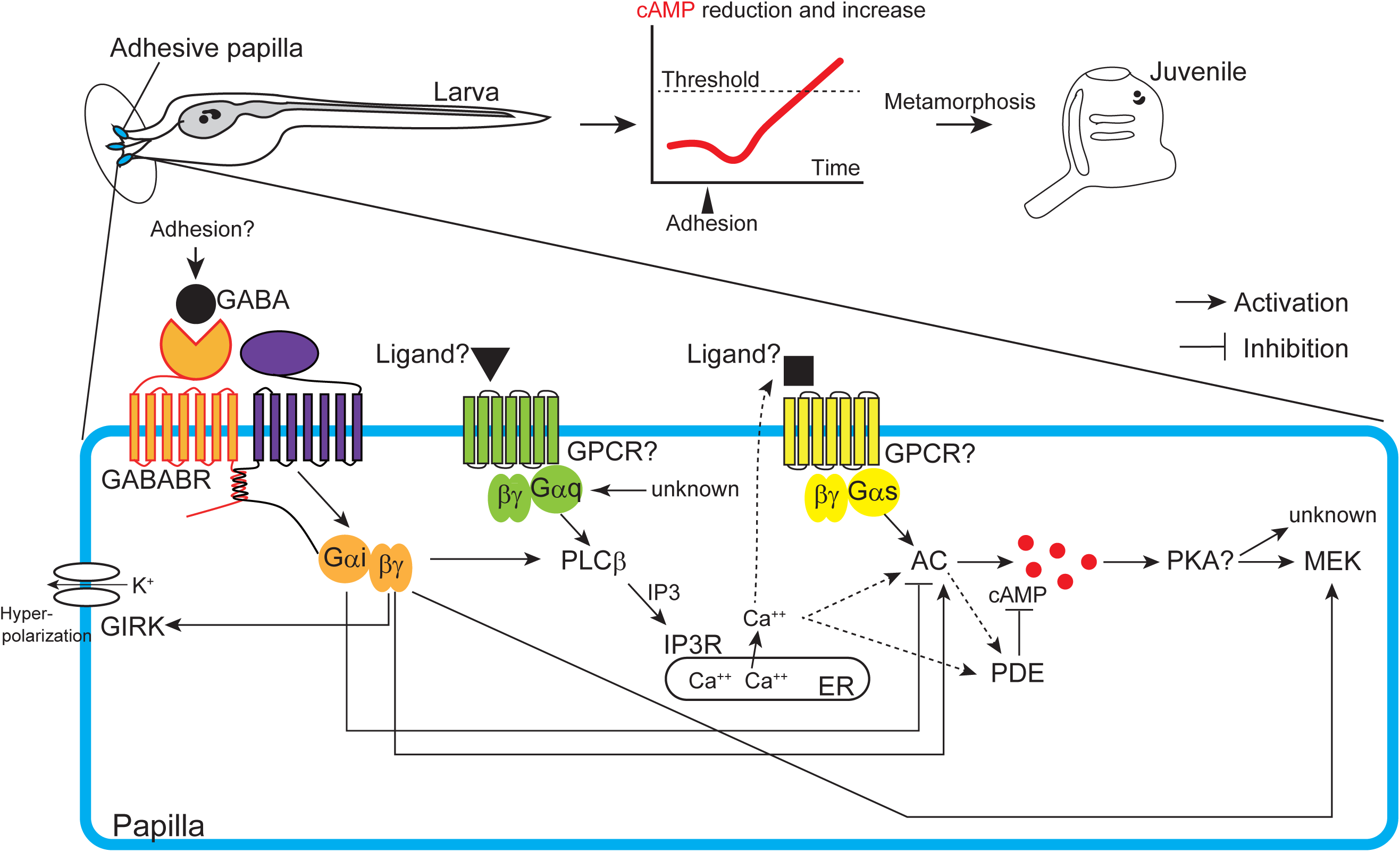
Schematic illustration of the signaling cascades for initiating *Ciona* metamorphosis. AC, adenylate cyclase; ER, endoplasmic reticulum; GPCR, G-protein coupled receptor; PKA, protein kinase A.

### Mechanisms measuring duration and strength of adhesions

One mystery of *Ciona* metamorphosis is that its initiation requires continuous adhesion with the adhesive organ for a certain period. This requirement is suggested for the faithful achievement of metamorphosis only when adhesion is firm enough to allow the transition out of the free-living larval stage. Our previous study suggested that approximately 30 min of adhesion is necessary^31^. A shorter duration of adhesion does not trigger metamorphosis. Larvae that experience short-term adhesion (such as detaching before 30 min) need to adhere again for 30 min to initiate metamorphosis, suggesting that the experience of temporal adhesion is erased. Our recent study showed that the force given to the papillae by adhesion also affects metamorphosis initiation: weaker force extends the time requirement^26^. Therefore, *Ciona* larvae likely possess a mechanism that somehow measures the duration and strength of adhesion and the ability to cancel transient excitations of the adhesive organ. Because fewer than 300 neurons constitute the larval nervous system of *Ciona*^62,63,83^, how the larva measures the strength and duration of adhesion with its simple nervous system is an important question to address the mechanisms of metamorphosis.

This study showed that cAMP is an essential second messenger molecule for triggering metamorphosis. Because PDE constitutively degrades cAMP, accumulation of this molecule requires persistent and/or strong activation of the metamorphic pathway that overcomes PDE activity. In other words, the duration and strength of adhesion could be converted into the quantity of cAMP, and when its quantity reaches a threshold, metamorphosis can be initiated. The time between the start of adhesion and metamorphosis initiation may be the period necessary to accumulate sufficient cAMP. This “cAMP timer” mechanism does not demand a complicated neuronal network system and could work in the simple nervous system of *Ciona*. Prior to triggering the cAMP timer, *Ciona* temporarily decreases cAMP quantity (Figure 3E and S4A). This reduction is also associated with metamorphosis initiation, suggesting that *Ciona* does not simply sense the cAMP increase. There are two plausible functions of this reduction. One is guaranteeing that the requirement of long-term adhesion will be met until a sufficient quantity of cAMP is accumulated, irrespective of the initial quantity of cAMP. The other one is multiplying the degree of cAMP increase. By reducing the initial value, larvae can increase the amount of this molecule several fold, even though the maximum quantity remains the same. Such an elevated difference in cAMP quantity may be adopted to ensure that the timing of metamorphosis initiation is appropriate using this ubiquitous molecule.

### G-protein signaling relay for metamorphosis

This study showed that three major G-proteins are involved in the initiation of metamorphosis (Figure 7). This complicated mechanism stands in contrast to the simple determination of metamorphosis initiation by cAMP quantity. What factor promoted the ancestor of ascidians to acquire this complicated signaling network of metamorphosis? We suspect that incorporating multiple components increases the chance of generating the crosstalk between molecules, providing robustness and flexibility to the system. Particularly, the involvement of Gi may have been important for achieving metamorphosis at the correct time and condition. GABA and Gi have inhibitory and excitatory functions^84,85^. *Ciona* larvae must ignore transient adhesion or stimulation on the papilla, which may occur frequently during swimming by colliding with an object or by strong water flows. At the same time, larvae must maintain sensing ability and start metamorphosis when they encounter an appropriate stimulus. Through the GIRK channel and Gαi, GABA can suppress the excitation and cAMP accumulation in the papilla. The GIRK channel could increase the excitation threshold of papilla neurons, allowing them to ignore weak stimuli. The transient reduction of cAMP at the onset of adhesion could be partly explained by the inhibitory function of Gαi on ACs. Not only Gi, but Gq and Gs are also necessary for this cAMP reduction. PDE1 is known to be activated by Ca^2+^/calmodulin^86^. Because PDE1 is abundantly expressed in the papillae (Figure S3B), the increase in Ca^2+^ upon adhesion by Gi and Gq could also trigger sudden cAMP reduction through PDE1. The contribution of Gs to cAMP reduction is surprising because the general role of this complex is to increase cAMP. However, it is known that an important target of the Gs pathway for the termination of cAMP synthesis is PDE^86^. The involvement of three G-protein pathways suggests that the acute cAMP reduction is a special event that is distinct from its constitutive degradation.

Together with its inhibitory function, GABA can activate the Gq pathway, which promotes metamorphosis through Gβγi. Group III AC and MEK could also be the targets of Gβγi. When adhesion is long enough, their activations somehow overcome the inhibitory functions of GABA. The GABA pathway is an ideal player that fulfills multiple requirements in the regulation of metamorphosis through its excitatory and inhibitory functions. We suspect that papillae themselves are the source of GABA for metamorphosis initiation. Because VIAAT/VGAT is not expressed in the papillae, the secretion of GABA from the papilla neurons requires an atypical mechanism. GABA is detected in the cell body of the neurons^23^. The release of cytosolic GABA may be triggered by the reception of mechanical stimuli using the TRP channel^26^. A VIAAT-independent cytosolic GABA release is observed in pancreatic beta cells^87^. Curiously, papilla neurons express transcription factor Islet^88^, which is homologous to the vertebrate transcription factor responsible for β cell function. The atypical secretion of GABA might depend on a transcription factor like Islet shared between *Ciona* papilla neurons and vertebrate beta cells.

A recent study^19^ provided several lines of evidence suggesting that papilla neurons (PNs) can serve as the sensors of several chemicals in addition to mechanical stimuli. This finding and our model could be mutually related because these chemicals could modify Ca^2+^ and cAMP production. G protein signaling allows *Ciona* to reflect various environmental stimuli to initiate metamorphosis either mechanically or chemically according to the situation.

### GPCRs for initiating metamorphosis and atypical Gαq activity

Future studies need to specify the mechanisms underlying Gαi, Gαq, and Gαs activation as well as the interaction between them more precisely. The activation of trimeric G-proteins relies on GPCRs, suggesting the presence of the receptors coupled with Gi, Gq, and Gs for metamorphosis initiation (Figure 7). Among them, Gi is likely to be coupled with GABABR; however, the expression of GABABR in the papillae is weak, and their interaction needs to be clarified in the future. We found that wild-type Gαq exhibits constitutive activity that does not require activation through adhesion. The constitutive Gαq activity requires GTP binding. If a GPCR owns this Gαq activation, its ligand should be supplied constitutively. Wild-type *Ciona* does not initiate metamorphosis without adhesion even though Gαq is expressed in the adhesive papillae, suggesting that the quantity of Gαq is strictly regulated to prevent autonomous metamorphosis in normal conditions.

We did not identify the GPCR coupled with Gs in the metamorphic mechanism. We suspect that GnRH receptors (GnRHRs) are strong candidates for this role. Our previous studies showed that GnRHs can induce metamorphosis^39,40^ as a downstream factor of GABA. Among the four genes encoding GnRHR proteins, *GnRHR1* and *GnRHR2* are expressed in the adhesive papillae^89^. These GnRHRs stimulate cAMP signaling, and the ligand GnRHs for GnRHR1 are also expressed in the papillae^89–91^. Therefore, GnRHs could be secreted from adhesive papillae as downstream molecules of GABA and stimulate the Gs pathway through GnRHR1. Future studies targeting the regulation of the secretion and reception of GnRH peptides after stimulation of papillae will improve our understanding of the signaling cascade conducting *Ciona* metamorphosis.

Understanding whether the G-protein signaling relay occurs in cell-autonomous or non-autonomous fashions is crucial for deepening our understanding of *Ciona* metamorphosis. Each papilla comprises approximately 20 cells, including four PNs. We showed that activation of Gq and Gs pathways in PNs is sufficient for initiating metamorphosis. This coincides with the previous reports^26,27^ that the loss of PNs by disrupting the POUIV transcription factor caused the complete arrest of metamorphosis. However, we do not think PNs are the only cells that activate G-proteins, because Ca^2+^ and cAMP imaging showed upregulation of fluorescence in the entire papillae^18,26^. A recent study also reports Ca^2+^ transient in another cell type than PNs^19^. Other papilla cells are likely responsible for activating G-proteins by directly responding to adhesion and by receiving signal input from adjacent cells, thereby enhancing the signal strength to accumulate a sufficient quantity of cAMP in the PNs to initiate metamorphosis. In this study, technical limitations prevented us from characterizing cells exhibiting increases in Ca^2+^ and cAMP; we need to address this question in future studies.

### Evolutionary implications

Sessile or benthic marine invertebrates lose locomotive activity when they undergo metamorphosis triggered by adhesion to a substratum^92^. These animals are suspected of having a system to control the adhesion state, allowing them to repeat attachment and detachment before meeting the appropriate conditions for metamorphosis. For example, barnacle larvae “walk” on the substratum before adhering firmly by secreting cement^93^. Like *Ciona*, the larvae of some sessile/benthic animals may have a system to initiate metamorphosis only when appropriate adhesion is provided, while erasing stimuli from transient and inappropriate adhesion. GABA serves as the metamorphosis inducer of some benthic invertebrates, including mollusks and echinoderms^94–96^. Moreover, GPCRs are implicated as the mediators of settlement and metamorphosis induction in hydrozoans, mollusks, and barnacles^94,97,98^. In the abalone and barnacle, the dual use of Gq and Gs pathways in metamorphosis has been suggested^99,100^. Supported by these shared features in the metamorphic mechanisms, our working hypothesis about the initiation of *Ciona* metamorphosis (Figure 7) will serve as a cue to elucidate how marine benthic invertebrates regulate their metamorphosis.

## Materials and Methods

### Animals

#### Ciona intestinalis

Type A wild types collected from Onagawa Bay (Miyagi, Japan) and Onahama Bay (Fukushima, Japan) were cultivated in closed colonies by the staff of the National BioResource Project, Japan. They were kept under a constant light condition to prevent gamete release. Eggs and sperm were collected surgically from gonadal ducts, and insemination was carried out in dishes. To prevent larvae from initiating metamorphosis, the tail’s posterior half was manually cut with a scalpel. Removing the tail prevents larvae from swimming efficiently, and these larvae are usually unable to adhere to a substrate. These adhesion-prevented larvae do not metamorphose, since adhesion is required to initiate metamorphosis. Larvae developed from dechorionated eggs were cultured on a 2% agar-coated dish after tail amputation to prevent metamorphosis. Because larvae stick to plastics, tail amputation is insufficient to avoid their adhesion to the culture dish.

### Pharmacological treatment

Larvae or larval anterior fragments isolated with a scalpel were administered overnight with 700 μM GABA (Fujifilm Wako #010-02441) or 1 mM theophylline (Sigma-Aldrich #T1633) dissolved in seawater, or 4 μM U0126 (Promega #V1121) dissolved in DMSO.

### Plasmids

The open reading frame (ORF) of Kaede was removed from pSPCiPC2Kaede by inverse PCR with PrimeStar GxL DNA polymerase (Takara-bio #R050). The ORFs of *Gαq*, *Gαs*, *Gαi*, and *PLCβ1/2/3* were PCR amplified using full-insert cDNAs^101^. These PCR fragments were fused with the In-Fusion HD Cloning kit (Clontech #639650). The region encoding XY linker^51^ was deleted from *PLCβ1/2/3* cDNA by inverse PCR to create a constitutively active form. The mutations were introduced by inverse PCR to create constitutive active or dominant-negative forms of *Gα* cDNAs. The introduced mutations are as follows: ca*Gαq*, corresponding to Q223L, abolishes GTPase activity^50^; ca*Gαs*, corresponding to Q227L, abolishes GTPase activity^55^; dn*Gαi*, corresponding to S47C^72,73^; and dn*Gαq*, corresponding to S54N^72^. The ORF of *bPAC*^59^ was PCR amplified and inserted between the *Bam*HI and *Eco*RI restriction sites of pSP-Kaede^102^ to create pSPbPAC. The PCR-amplified *PC2 cis*-element^54^ was inserted into the *Bam*HI site of pSPbPAC. The plasmid DNAs were linearized by a restriction enzyme and purified with the Qiaquick Gel Extraction Kit (Qiagen #28706) before microinjection. The ORF of *Pink Flamindo*^60^ was PCR amplified and inserted into the *Eco*RV restriction sites of pBS-HTB^103,104^ to create pHTBPinkFlamindo. The plasmids pHTBGCaMP8^26^ and pHTBPinkFlamindo were linearized using *Xho*I for subsequent *in vitro* synthesis of *GCaMP8* and *Pink Flamindo* mRNA, respectively. *GCaMP8* mRNA was synthesized with the MEGAscript T3 kit (Thermo Fisher Scientific #AM1338), the Poly (A) Tailing kit (Thermo Fisher Scientific #AM1350), and the Cap structure analog (New England Biolabs #S1404). Pink Flamindo mRNA was synthesized with the mMESSAGE mMACHINE T3 Transcription kit (Thermo Fisher Scientific #AM1348).

### Microinjection

Unfertilized eggs were dechorionated in sterilized seawater containing 1% sodium thioglycolate (Fujifilm Wako #590-11762) and 0.05% actinase E (Kaken Pharmaceutical #650164) as previously described^105^. The microinjection solution included 2 mg/ml of Fast Green FCF (Fujifilm Wako #061-00031), 0.5-1.0 mM of morpholino oligonucleotide(s) (MOs)^106^, 5 ng/ul of plasmids, and/or 1 mg/ml of *GCaMP8* mRNA for imaging^26^. For cAMP imaging, 1 mg/ml *Pink Flamindo* mRNA was dissolved in water without Fast Green, since this chemical emits fluorescence. Microinjected unfertilized eggs were inseminated in a gelatin-coated plastic dish. After the seawater was exchanged to remove excess sperm, the fertilized eggs were cultured at 18 °C or 20 °C overnight until the larval stage. MOs are listed below. Standard control MO, 5’- CCTCTTACCTCAGTTACAATTTATA-3’; *GAD* ATG MO^39^, 5’-ACCTCCAAGCCGATTGTTTCTGCAT-3’; *GABABR1* ATG MO^39^, 5’- GCTTACGACTTTACATAACCTTACA-3’; *Gαq* ATG MO, 5’- GGCATATTTGTGACTATAATGACG-3’; *Gαs* ATG MO, 5’- AAAGCAACCCATTGGCATTATCGAC-3’; *PLCβ1/2/3* splicing MO, 5’- GTGTTACTTACGCTTTCTCTA-3’; *PLCβ4* splicing MO, 5’- AACCACCAACCACCAACCTTTTG-3’; *IP3R* splicing MO, 5’- AATGATGGTTTAAAATTGCCACCTG-3’; *Gαi* ATG MO, 5’- GTGGAGACTGTGCAACCCATGATTC-3’; *dvGai_Chr2* ATG MO, 5’- CCATCTTGAGTAATCCAGGCTTTTA-3’; *dvGai_Chr4* ATG MO, 5’- CATGGTCAGCGGTTTACAAAGTATT-3’; *GIRK channel* ATG MO, 5’- TCTGCTGGTTCAGTAATAGACATAG-3’.

### Photographing and imaging

Photographs were taken with an AxioImager Z1 and AxioObserver Z1 (Carl Zeiss). The images were treated with AxioVision Rel.4.6 or Zen (Carl Zeiss) and Photoshop 2021 (Adobe) software. Imaging of the Ca^2+^ transient was carried out according to previous reports^18,26^. The tails of *GCaMP8* mRNA-injected larvae were removed by a scalpel at 15-17 hours after fertilization (hpf). At 30-40 hpf, the larvae were mounted onto a plastic dish under an M165FC stereo microscope (Leica Microsystems), and their adhesive papillae were stimulated by a glass rod using a three-axis coarse manipulator M-152 (Narishige Japan). Immediately after the tip of the rod had contact with the papillae, visible light was shut off and the fluorescence of GCaMP8 was taken with a DFC 310FX digital camera (Leica), Las version 4.12 (Leica), and the movie-capturing function of Windows 10. Images were treated with Photoshop 2021.

Imaging of cAMP was taken by confocal laser scanning microscopy (FV1000, Olympus). The movies were treated with ImageJ (National Institutes of Health). The Pink Flamindo mRNA-injected larvae were immobilized on Poly L lysine-coated glass-bottom dishes at 20-21 hpf, and their adhesive papillae were stimulated around 25 hpf. For the stimulation, an electric manipulator MM-89 (Narishige) and hydraulic micromanipulator MMO-202ND (Narishige) were used.

### RNA sequencing

Larval anterior portions, including adhesive papillae and the other trunk region, were separately isolated by a scalpel around 22-25 hpf at 18°C and collected in Isogen (Nippon Gene #317-02503) soon after isolation. To identify GABA-responsive genes, half of the tail was amputated from larvae around 21-24 hpf, followed by GABA administration for 2 hours before treatment with Isogen. In each experiment, approximately 50 fragments were collected, and two biological replicates were taken. Total RNA was isolated according to the manufacturer’s instructions. Ribosomal RNA was depleted from the total RNA using the NEBNext rRNA Depletion kit (New England Biolabs #E6310), followed by the conversion of the remaining RNA into an Illumina sequencing library using the NEBNext Ultra Directional RNA Library Prep kit (New England Biolabs #E7760). Following library preparation, validation was performed using the Bioanalyzer system (Agilent Technologies) to assess size distribution and concentration. Subsequently, sequencing was carried out on the NextSeq 500 platform (Illumina) employing the paired-end 36-base read option. Reads were mapped on the *Ciona intestinalis* Type A reference genome (HT model; http://ghost.zool.kyoto-u.ac.jp/default_ht.html) and quantified using CLC Genomic Workbench version 22.0 (Qiagen). RNA-seq data sets were deposited in the NCBI Gene Expression Omnibus (GEO) under accession numbers SAMN40712866 to SAMN40712873, which will appear online upon publication of this paper.

The read counts were normalized by calculating the number of reads per kilobase per million (RPKM) for each transcript in each sample using CLC Genomic Workbench version 22.0 (Qiagen). Eight candidate genes showing a minimum 1.5-fold increase following GABA administration, along with notably higher expression in the papillae in comparison to the trunk (an approximately 4-fold increase in a sample), were chosen for quantitative PCR analysis to assess their response to GABA.

### Quantitative PCR

Total RNA was isolated from the anterior tips of larvae treated with the chemical using Isogen (Nippon Gene), following the manufacturer’s instructions. After isopropanol extraction, genomic DNA removal and reverse transcription were carried out using the PrimeScript™ RT reagent kit with gDNA Eraser (Takara Bio #RR047). Quantitative RT-PCR (qRT-PCR) was done using a SYBR Premix Ex TaqII Dimer Eraser (Takara Bio #RR091) and a Thermal Cycler Dice Real Time System III (Takara Bio). The gene encoding elongation factor 1α (*EF1α*) was used to normalize RNA quantity according to a previous study^104^. qPCR primers are listed below. KY21.Chr5.240, 5’- TCTTCTCAAAGTTGCACATTCC-3’ and 5’-CAGCAGCAACCAAACGATAAAC-3’; KY21.Chr8.489, 5’- CAATGCAACTTTGACTGCATAC-3’ and 5’- TCCAAACTGCATTCCACATATC-3’; *EF1α*, 5’-CATGTCACGGACAGCGAAACG- 3’ and 5’- CAATGTGTGTTGAGGCATTCCAAG-3’.

### Statistical analysis

Differences between conditions were evaluated by Fisher’s exact test. Statistical analyses and most graph visualizations were conducted utilizing R x64.4.1.2 and RStudio software.

### Phylogenetic analysis

We performed a tBlastn search^107^ against the HT gene model of *Ciona*^44^ using the amino acid sequences of human PDE, PLCβ, and *Ciona* Gα proteins as queries, followed by a reciprocal Blast search. We aligned the amino acid sequences from the HDc domain of PDE, the EF-hand_like, PLCXc, PLCYc, and C2 domains of PLCβ, and the whole open reading frame of Gα proteins using the M-COFFEE program^108,109^. After the removal of unnecessary residues, maximum likelihood trees were constructed using MEGA software version 11^110^, employing the WAG amino acid substitution matrix^111^. The trees were assessed with 1,000 bootstrap pseudoreplicates.

## Acknowledgements

We would like to express our earnest gratitude to Drs. Kazuki Horikawa, Atsuo Nishino, Ryusuke Niwa, and Takahiro Yamashita for their kind material provision, helpful comments, and general support of our study. We thank Dr. Kogiku Shiba for instructing us with the pharmacological analyses. We also thank Dr. Masafumi Muratani and members of i-Laboratory at the University of Tsukuba for their support for RNA sequencing. We are grateful to the past and present members of the Shimoda Marine Research Center at the University of Tsukuba for their contributions to the initial step of this study and the maintenance of the animals. We thank Drs. Shigeki Fujiwara, Manabu Yoshida, Yutaka Satou, and all members of the Department of Zoology, Kyoto University, the Misaki Marine Biological Station, the University of Tokyo, the Maizuru Fishery Research Station of Kyoto University, and the National BioResource Project (NBRP) for the cultivation and provision of *Ciona* adults and experimental materials. This study was supported by grants from the Japan Society for the Promotion of Science to K.H. (21H00440, 23H04717), T.H. (19H03204, 21K19249, 21H05239), and Y.S. (19H03262). Y.S. was further supported by a Takeda Bioscience Research Grant. T.H. was supported by the Collaborative Research in Computational Neuroscience program (CRCNS2021). N.M.T was supported by JST SPRING (JPMJSP2123).

## Author Contributions

A.H., N.M.T., A.O., Y.W., and Y.S. performed the experiments. Y.S. designed the experiments. T.H., M.H., and A.S. performed RNA-seq data and phylogenetic analyses. H.S., T.H., and K.H. supervised the work. Y.S. wrote the manuscript with consultations by A.H., N.M.T., and K.H. All authors contributed to the manuscript preparation.

## Figure legends for Supplementary Figures

**Supplementary Figure S1.**
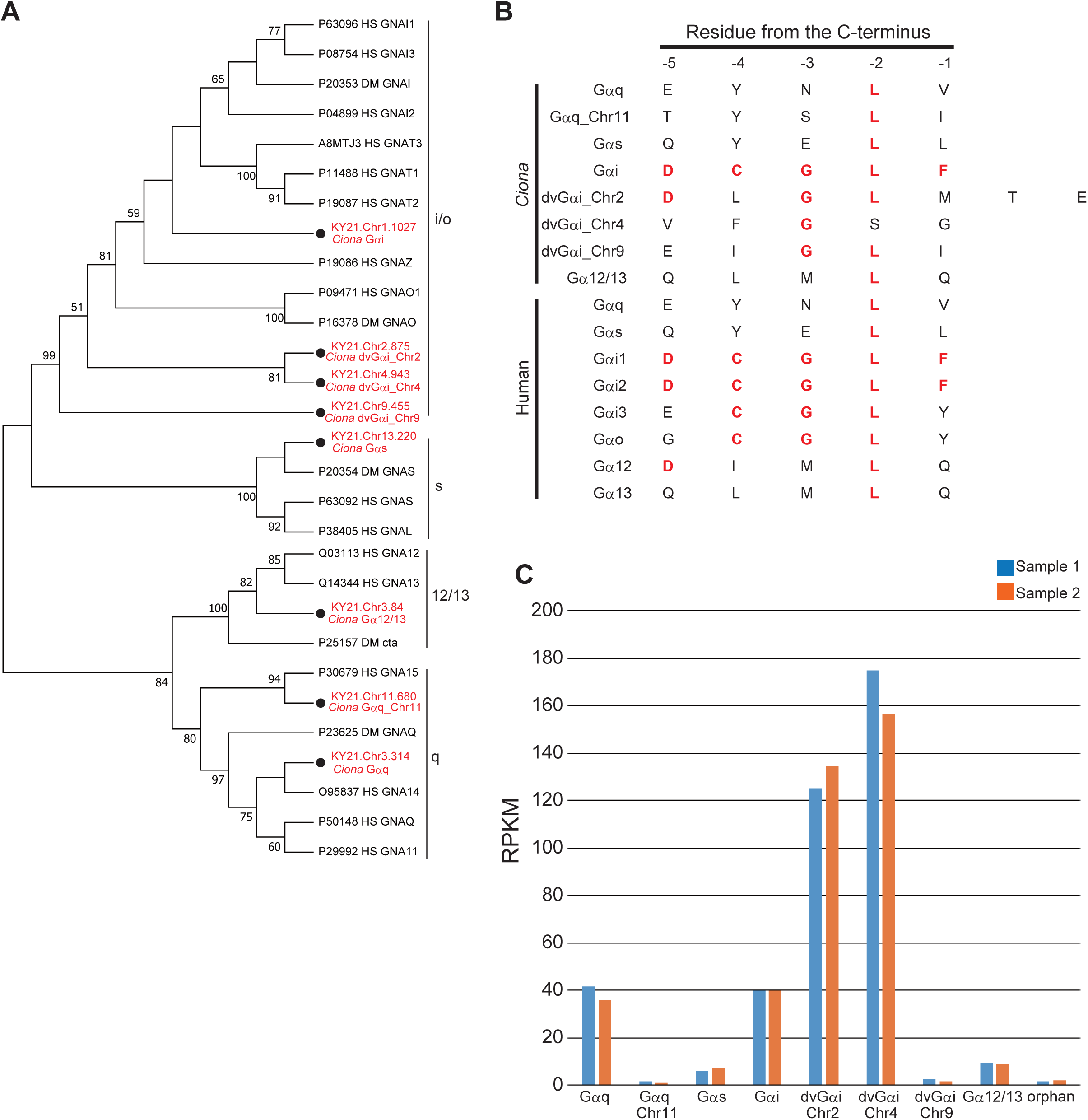
*Ciona* genes encoding the G alpha subunit. (A) Phylogenetic relationships between Gα proteins, as revealed by the maximum likelihood method. *Ciona* proteins are highlighted in red. (B) Comparison of the five C-terminal amino acid sequences of Gα proteins that are shown to be essential for their functions, including two additional amino acids for dvGαi_Chr2. Residues conserved with human Gαi1 are shown in red. (C) Expression profiles of Gα genes, as revealed by RNA-seq of the papilla region. RPKM, reads per kilobase of exon per million mapped reads; “orphan”, the gene represented by the model KY21.Chr8.580. See also Table S1.

**Supplementary Figure S2.**
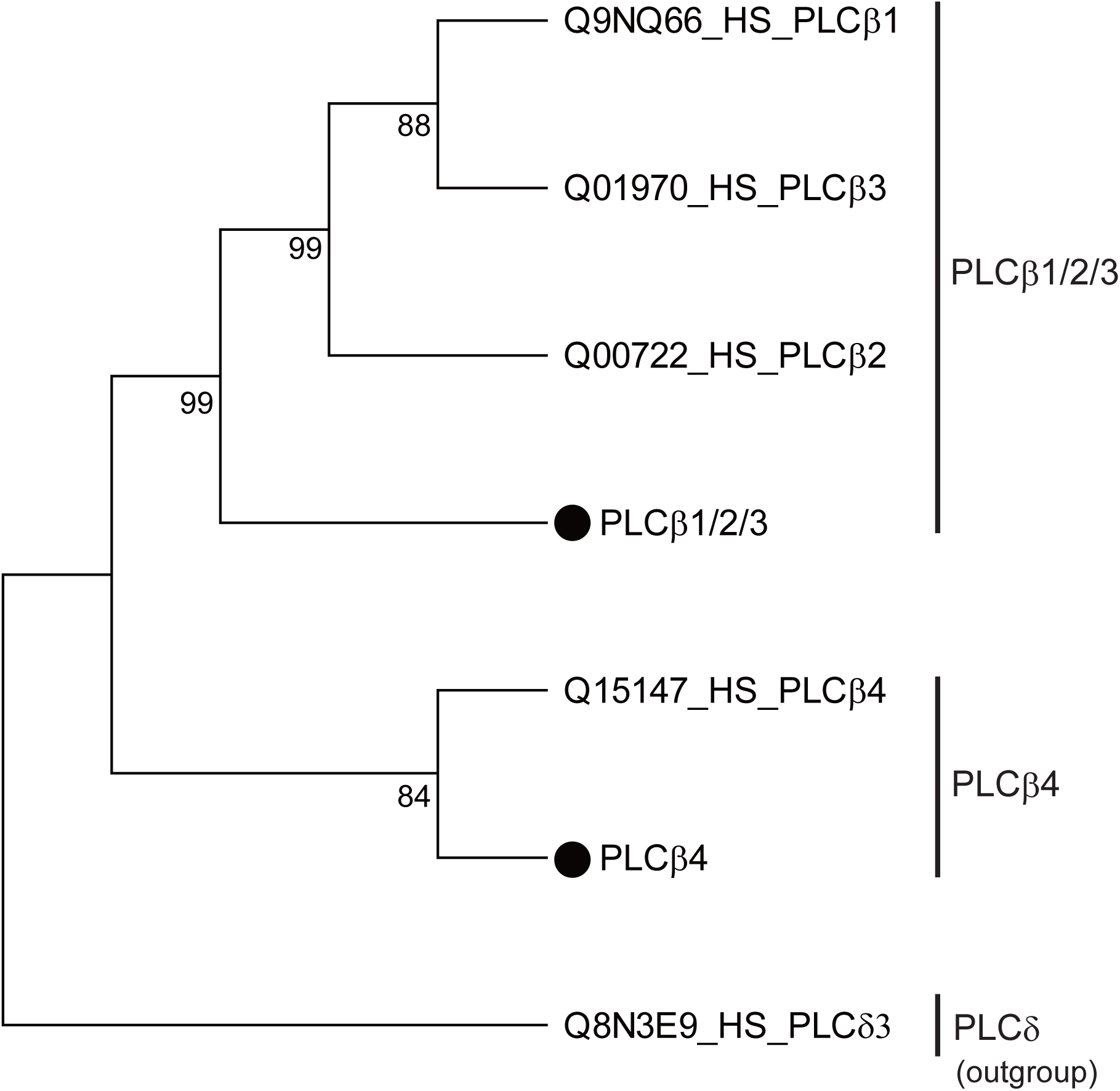
*Ciona* phospholipase β proteins. Phylogenetic relationships between PLCβ proteins, as revealed by the maximum likelihood method. The number beside each branch indicates the percentage of times that a node was supported in 1,000 bootstrap pseudoreplications. Scores equal to or exceeding 50% are shown. *Ciona* PLCβs are indicated by black circles. Human homologs are indicated by accession numbers in the Uniprot database (https://www.uniprot.org) and “HS” for *Homo sapiens*.

**Supplementary Figure S3.**
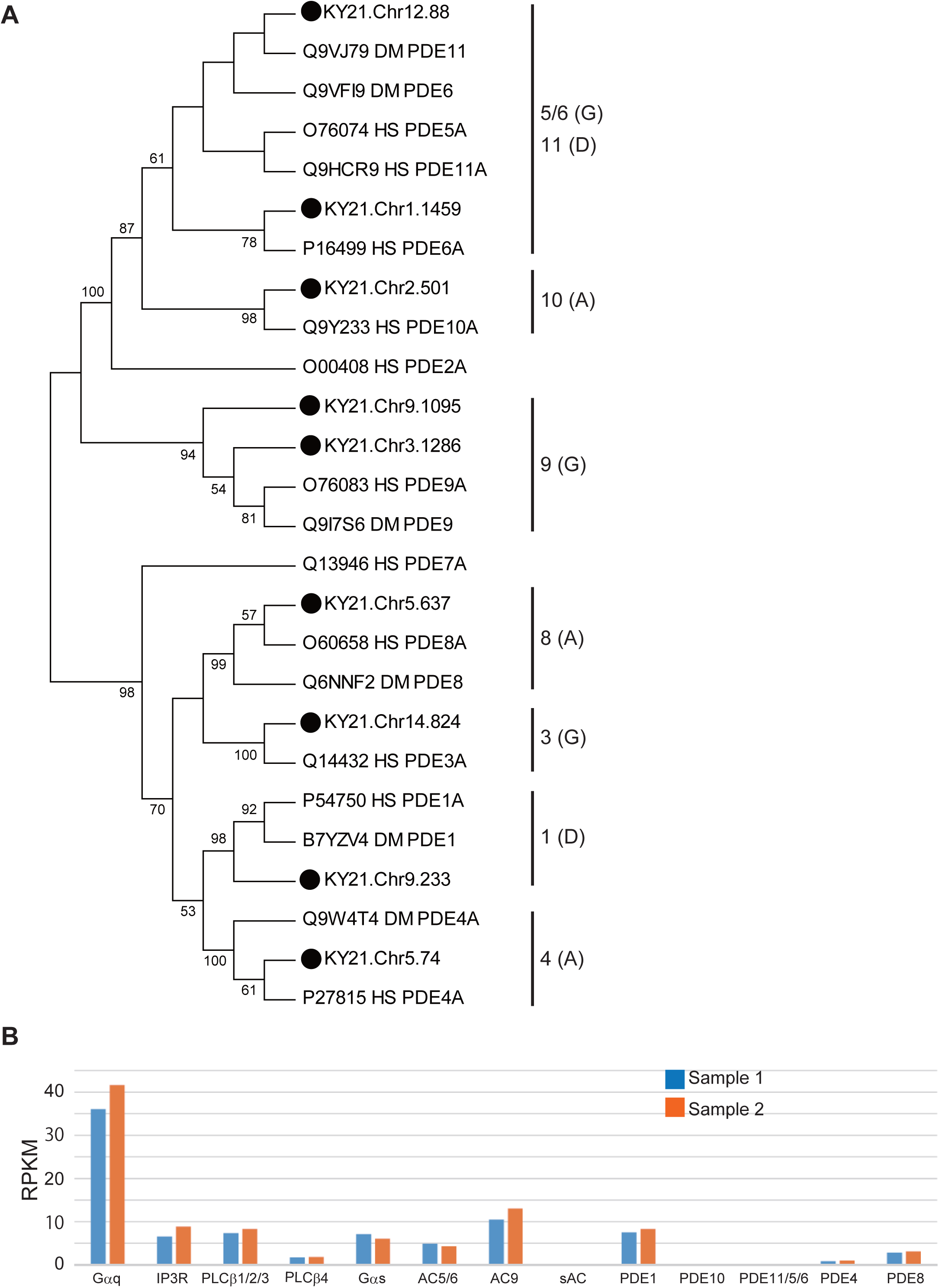
*Ciona* phosphodiesterase repertoire and expression of Gq and Gs pathway genes in the adhesive papillae. (A) Phylogenetic relationships between PDEs. Black circles and gene models indicate *Ciona* protein models in the Ghost database (http://ghost.zool.kyoto-u.ac.jp/default_ht.html). The numbers and letters in parentheses indicate the PDE groups and their suspected substrates. A, cAMP; G, cGMP; D, dual substrate. Human and insect proteins are shown by Uniprot (https://www.uniprot.org) accession numbers, as well as “HS” for *Homo sapiens* and “DM” for *Drosophila melanogaster* proteins. (B) Expression profiles of the genes related to Gq and Gs pathways, as revealed by RNA-seq of the papilla region. RPKM, reads per kilobase of exon per million mapped reads. See also Table S1.

**Supplementary Figure S4.**
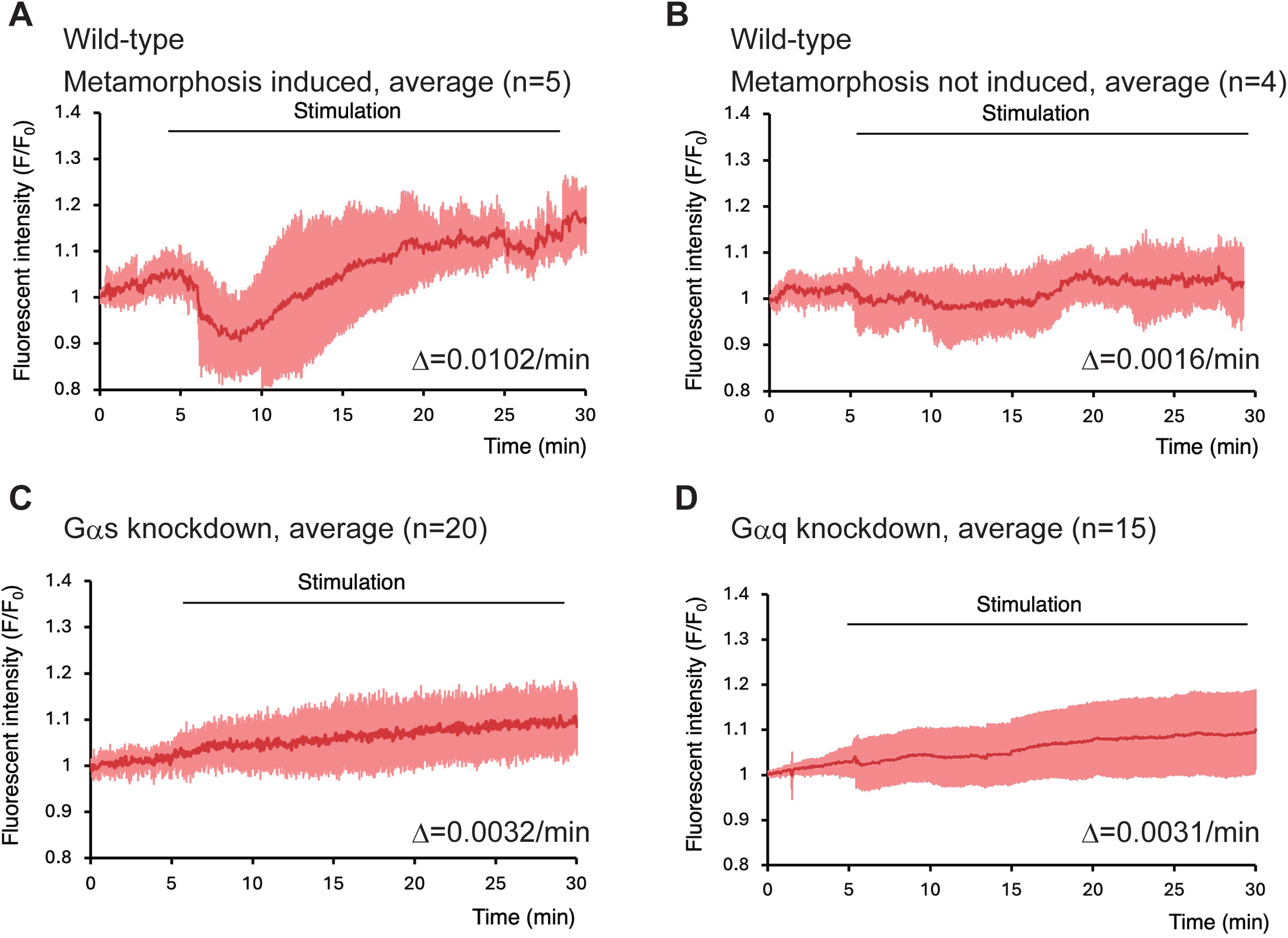
cAMP increase in the papillae is coupled with metamorphosis initiation. (A) Measurement of cAMP concentration in the larva that initiated metamorphosis upon stimulation of the papillae, as indicated by the fluorescence of Pink Flamindo. The red and pink graphs represent the averaged Pink Flamindo fluorescent intensity and the standard deviations, respectively. The number in the parentheses indicates the number of examined larvae. The red graph after the initiation of fluorescent intensity increase is approximated to the linear graph to calculate its slope, which is shown as Δ. (B) Measurement of cAMP concentration in the larva that did not initiate metamorphosis upon stimulation of the papillae, as indicated by the fluorescence of Pink Flamindo. The red graph is approximated to the linear graph to calculate its slope, which is shown as Δ. Note that this larva was not subjected to any experimental perturbation of genes. (C) Measurement of cAMP concentration in the *Gαs* knockdown larvae. (D) Measurement of cAMP concentration in the *Gαq* knockdown larvae.

**Supplementary Figure S5.**
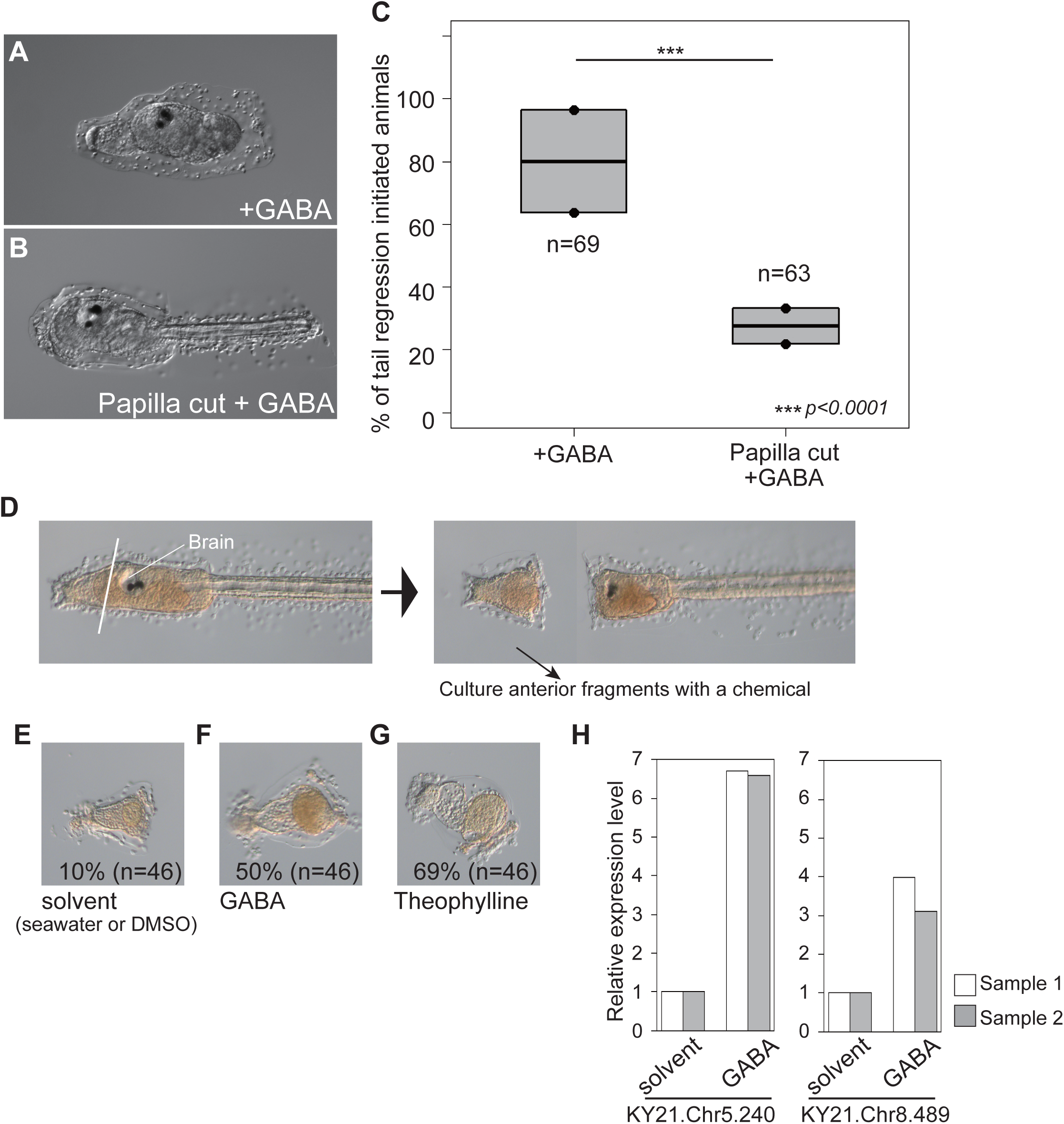
GABA stimulates the adhesive papillae to initiate metamorphosis. (A) A GABA-treated animal initiated metamorphosis. (B) A papilla-amputated larva treated with GABA. Metamorphosis was not initiated. (C) Effect of papilla amputation on GABA treatment. (D-H) The sensory vesicle is not required for the anterior region to respond to GABA. (D) Schematic of the experiment. (E) A control anterior fragment at 2 dpf. (F) Anterior fragment treated with GABA. The percentage indicates the proportion of the fragments exhibiting the morphological signature of metamorphosis. (G) Anterior fragment treated with theophylline. (H) Expression levels of two GABA-responsive genes in the control- and GABA-treated anterior fragments, as revealed by quantitative RT-PCR.

**Supplementary Figure S6.**
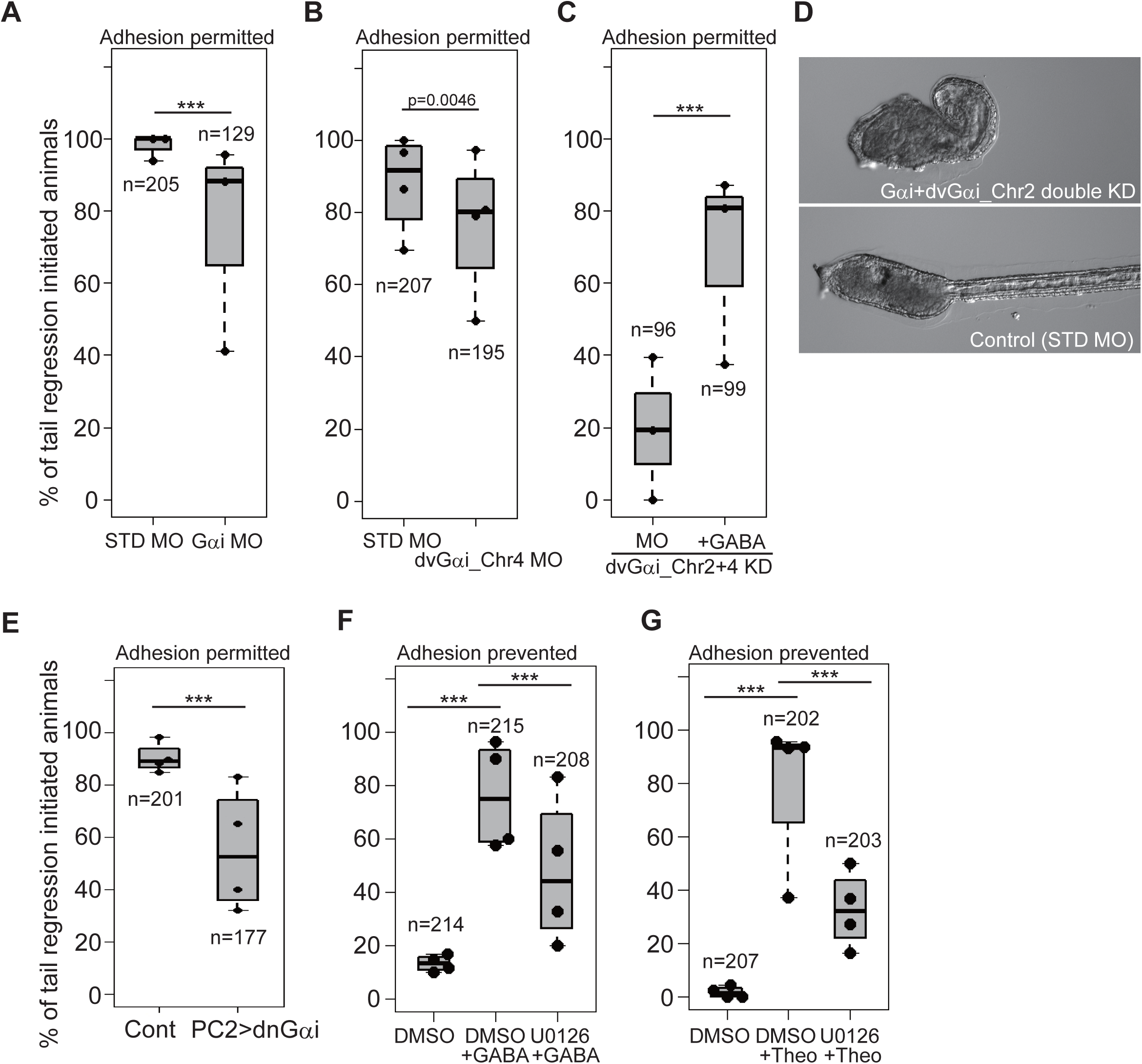
Gi is required for metamorphosis. (A) Effects of *Gαi* knockdown. (B) Effects of *dvGαi_Chr4* knockdown. (C) Effects of simultaneous knockdown of *dvGαi_Chr2* and *dvGαi_Chr4*. (D) Effects of simultaneous knockdown of *Gαi* and *dvGαi_Chr2* on morphology. (E) Effects of dn*Gαi* overexpression. (F) Effects of U0126 on GABA treatment. (G) Effects of U0126 on theophylline treatment.

**Supplementary Figure S7.**
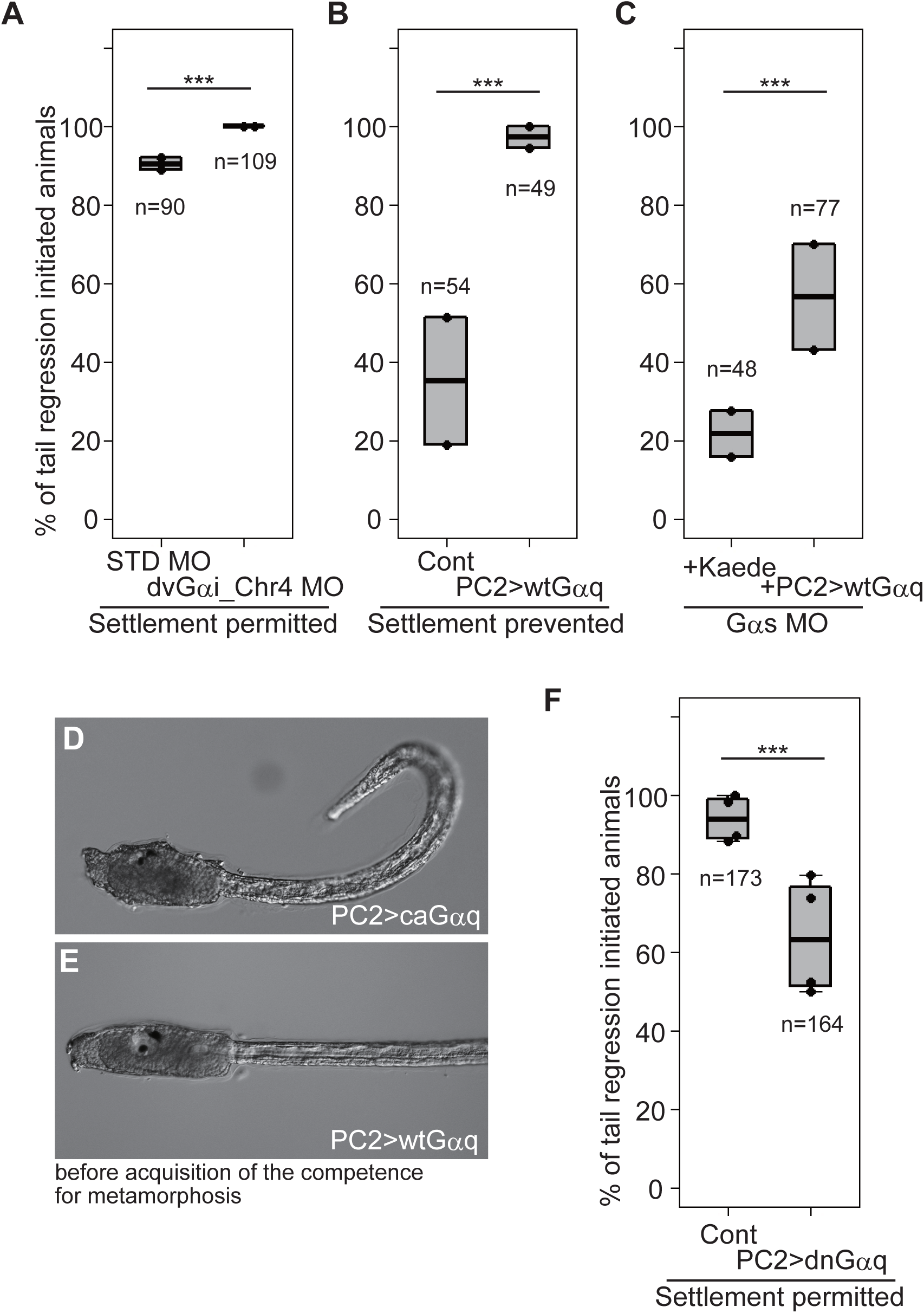
Autonomous activity of wild-type Gαq. (A) Effect of wt*Gαq* overexpression in the adhesion-permitted condition. (B) Effect of wt*Gαq* overexpression in the adhesion-prevented condition. (C) Effect of wt*Gαq* overexpression on *Gαs* knockdown. (D) Morphology of ca*Gαq*-overexpressed larva at 1 dpf (before initiation of metamorphosis). (E) Morphology of wt*Gαq*-overexpressed larva at 1 dpf. (F) Effect of dn*Gαq* overexpression in the adhesion-permitted condition.

**Supplementary Table S1.**
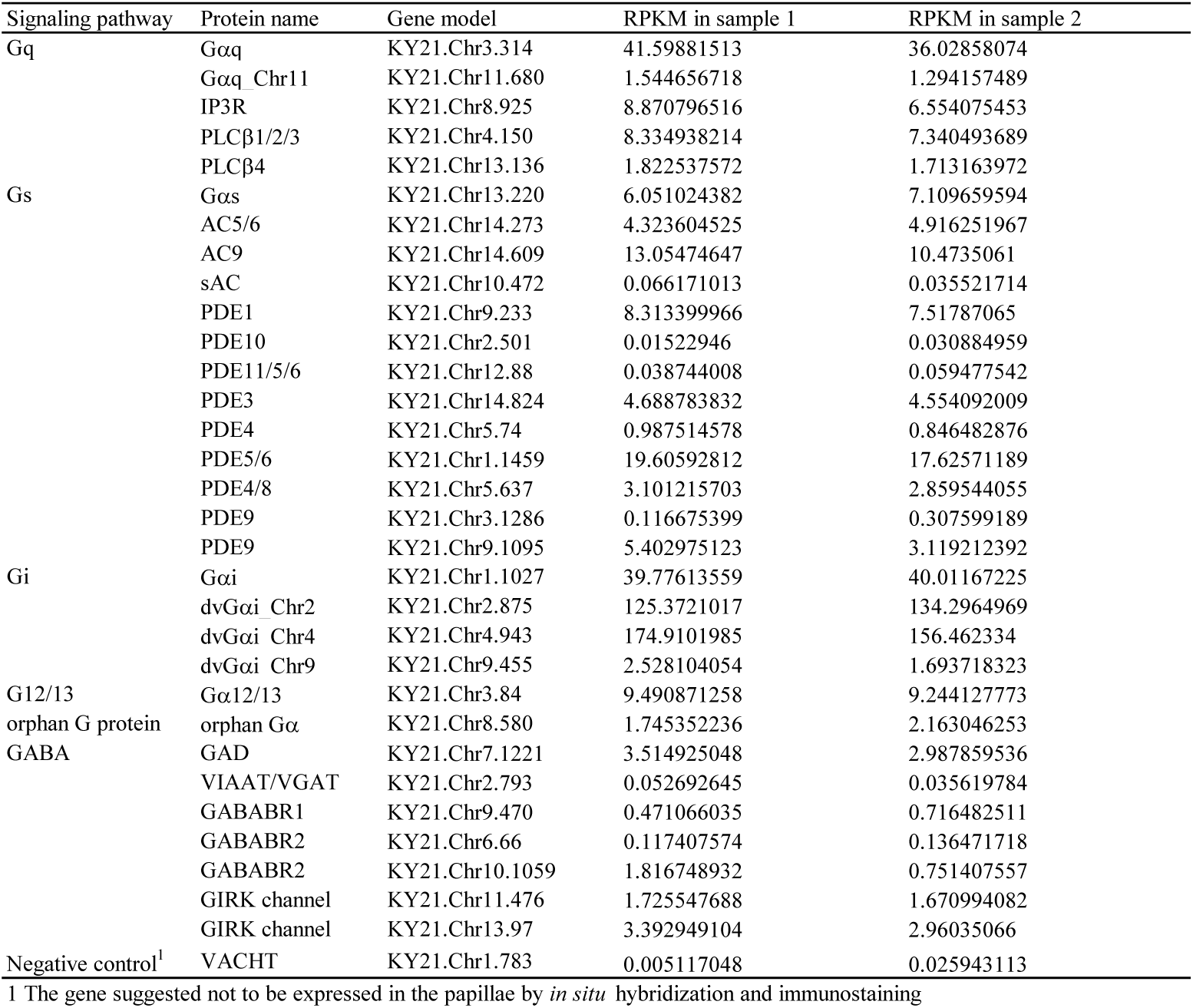
Expression levels of genes related to G-protein signaling during metamorphosis, as revealed by RNA-seq of the larval anterior tip including papillae.

**Supplementary Table S2.** List of candidate genes upregulated by GABA.

| KY21 gene model | Human homolog | <sup>1</sup>GABA vs Control<br>Sample 1 | <sup>1</sup>GABA vs Control<br>Sample 2 | <sup>2</sup>Papillae vs Trunk<br>Sample 3 | <sup>2</sup>Papillae vs Trunk<br>Sample 4 |
| --- | --- | --- | --- | --- | --- |
| KY21.Chr1.1223 |  | 2.33 | 1.97 | Not determined | Not determined |
| KY21.Chr2.1293 |  | 2.07 | 3.09 | Not determined | Not determined |
| KY21.Chr2.219 |  | 1.78 | 1.8 | Not determined | Not determined |
| KY21.Chr2.1069 | ASNS: asparagine synthetase (glutamine-hydrolyzing) | 6.37 | 11.6 | 22.05 | 49.41 |
| KY21.Chr5.240 | Serine peptidase inhibitor Kunitz type 3 | 1.79 | 2.11 | 12.57 | 14.39 |
| KY21.Chr8.1171 |  | 1.56 | 1.52 | 11.08 | 10.04 |
| KY21.Chr3.438 |  | 2.38 | 1.91 | 8.61 | 9.16 |
| KY21.Chr11.358 |  | 3.57 | 3.39 | 4.69 | 1.2 |
| KY21.Chr9.130 | CTH: cystathionine gamma-lyase | 2.16 | 2.75 | 4.68 | 3.26 |
| KY21.Chr9.1057 | ASS1: argininosuccinate synthase 1 | 3.69 | 6.23 | 4.5 | 3.31 |
| KY21.Chr3.1611 |  | 1.69 | 1.54 | 4.48 | 7.14 |
| KY21.Chr1.732 |  | 2.34 | 1.66 | 4.37 | 2.74 |
| KY21.Chr8.489 | AIFM2: apoptosis-inducing factor, mitochondria-associated 2 | 1.85 | 2.05 | 4.01 | 1.77 |
| KY21.Chr7.224 | B4GALT1: beta-1,4-galactosyltransferase 1 | 2.43 | 1.51 | 3.99 | 4.16 |

1 The RPKM in the GABA-treated sample was divided by the RPKM in the solvent (seawater)-treated sample to calculate the scores.
2 The RPKM in the papilla region was divided by the RPKM in the remaining trunk region to calculate the scores.

